# Carbon nanoparticle exposure strengthens water-relation parameters by stimulating abscisic acid pathway and aquaporins genes in rice

**DOI:** 10.1101/2024.02.11.579789

**Authors:** Aman Kumar, Manasa S Lekshmi, Jyotiprabha Kashyap, Sikha Mandal, Gayatri Mishra, Jnanendra Rath, Gyana Ranjan Rout, Kishore CS Panigrahi, Madhusmita Panigrahy

## Abstract

Mechanism of action and molecular basis of positive growth effects including yield increase due to carbon nanoparticle (CNP) treatment in rice plants is dissected here. CNP at 500 -750 µg/mL were found to be the optimum dosages showing best seedling growth. CNP treatment resulted increase in stomata size, gaseous exchange and water use efficiency along with decrease in stomata frequency, relative humidity, internal CO_2_ concentration. CNP treatment exerted cold tolerance in seedlings and water stress tolerance in reproductive stage. CNP-coupled with water uptake was found to be endocytosis mediated, although CNP uptake was not affected by endocytosis inhibitor application in roots. Genomic analysis resulted major involvement of ABA pathway and stomata size and frequency genes in *Arabidopsis* and rice. Elevated endogenous ABA in rice seedlings and flag leaves along with increased expression of ABA biosynthetic genes in *Arabidopsis* and rice *AtNCED3*, *AtNCED6*, *OsNCED1* confirmed increased ABA synthesis. Negative regulators of ABA pathway, *OsSNRK2* down-regulation and up-regulation of stomagen (*OsEPFL9*) reconfirmed ABA’s involvement. CNP treatment resulted water stress tolerance by maintaining lower stomatal conductance, transpiration rate and higher relative water content. Increased ABA (*OsSNRK1*, *OsSNRK2*) and aquaporin (*OsPIP2-5*) genes’ expressions could explain the better water stress tolerance in rice plants treated with CNP. Altogether, due to thermomorphogenesis, down-regulation of Phytochrome B resulted altered the ABA pathway and stomatal distribution with size. These changes resulted improved water relation parameters and WUE showing improvement in yield. Detailed mechanism of action of CNP in abiotic stress tolerance can be exploited in in nano-agriculture.

## Introduction

Use of nanomaterials for improvement in agriculture is known as nano agriculture and the technologies exploited for these purposes are together called nanotechnology. Nano agriculture in this emerging era has proved as a potential tool to solve several problems faced in agriculture (Mazeed et al., 2020). The multifaceted solutions offered by nano agriculture include avoiding excessive use of pesticides in crops, increased growth and yield of crops, targeted delivery of DNA at the site of application, improved nutrient use efficiency, detection of diseases by the use of nanosensors etc. (Rani et al., 2020). Use of nano materials greatly effects the plant species through different pathways and influences their effectiveness. Hence, the method of their application is very critical to meet the expectations (Alejandro, 2017). Equally important is to understand how the nanomaterials are absorbed, transported, internalized and brings its effect in the plant species in which they are applied. As it strengthens the fundamentals of the mechanisms involved and clarifies the issues related to wide acceptance of the method. Recent reports have also started to explore the mechanisms underlaying and the working principles of the nano materials inside plants of use. Studies have attempted to address the molecular mechanisms of nano toxicity using differential protein expression profiling, genomics approaches, microRNA analysis and nanotoxicoproteomics approaches (Jha et al. 2016). Studies have shown that in addition to the internal use of nanomaterials, their external application in the growth medium can also induce growth effects without toxicity issues. External application methods of engineered or non-engineered nanomaterials include foliar spray, hydroponics, seed priming, and in soil (Zhao et al., 2020). External application of multi walled carbon nanotubes (MWCNT) in the growth medium enhanced seed germination and growth by exhilarating expression of several aquaporins, plasma membrane intrinsic proteins and tonoplast intrinsic proteins (Lahiani et al., 2013). In our previous report, genetic mechanisms associated with external application carbon nanoparticle (CNP) in the growth medium of *Arabidopsis* resulted accelerated flowering, photoperiod dependence and shade avoidance (SAR)-like effects with down-regulation of red-light photoreceptor Phytochrome B’s transcript levels (Kumar et al., 2018). This was probably the first report elucidating links of the mechanisms of CNP with another physiological syndrome such as SAR and light signalling. Nanoparticle exposure have also been greatly exploited to attain robust growth and yield in different plant species (Zhao et al., 2020). Our effort to test the effect of CNP external application in the rice pots resulted improvement in growth and yield of rice plants (Panigrahy et al., 2021). The analyses of underlaying mechanisms revealed hike of plant internal temperature by 0.5°C ± 0.1°C due to the presence of CNP which could mimic the situation of shade and stimulated SAR in rice plants. Several recent studies have also successfully attempted to attain abiotic and biotic stress tolerance through external application of nanomaterials in plants (Ali et al. 2021, Majeed et al., 2020). An et al. have shown salinity tolerance in cotton by application of cerium oxide nanoparticles in the germinating medium (An et al., 2020). Hatami et al. showed improved drought tolerance in *Hyoscyamus* seedlings due to 14-days exposure to SWCNT under different levels drought induced by polyethylene glycol (Hitami et al., 2017). Despite so many analyses and advantages obtained from nanomaterials application, detailed study including natural environmental conditions covering full life cycle of a plant and addressing the nutritional quality of the grains obtained are scanty. Effect of cerium oxide nanoparticles on the rice grain quality was shown by Rico et al., (2013). Beneficial effects of copper oxide nanoparticles in rice at filed level with their effect on grain was shown by Deng et al., (2022). Our previous results demonstrated that CNP exposure in rice pots improved the nutritional quality of the rice grains obtained from the CNP-treated plants. In the present study we have investigated the molecular basis of action of CNP to exert the growth effects in rice. We have also accessed different abiotic stresses to find which abiotic stress tolerance CNP can induce in rice plants. Further, molecular, genetic as well as genomic analyses have been attempted to answer the basis of drought tolerance in rice plants that we obtained due to CNP exposure.

## Result

### Dosage dependence effect of CNP on the rice seedlings’ phenotypic growth

CNP had induced several positive growth effects in rice plants (Panigrahy et al., 2021). Hence, it was necessary to determine the ineffective, optimum and maximum concentrations of CNP to show positive growth effects. Study of seedling shoot length (SL), root length (RL), rootlet number (RLN) and nodal roots (NR) at 7 different CNP concentrations (ranging from 200 µg/ml to 1500 µg/ml) (Figure 1, Suppl Figure 1) showed that SL, RL and RLN tend to increase after at least 350 µg/ml indicating positive growth effects. SL and RL showed highest increase between 500 and 750 µg/ml after which they started to decline (Figure 1A). RLN also was highest number at 750 µg/ml (Figure 1).

**Figure 1.**
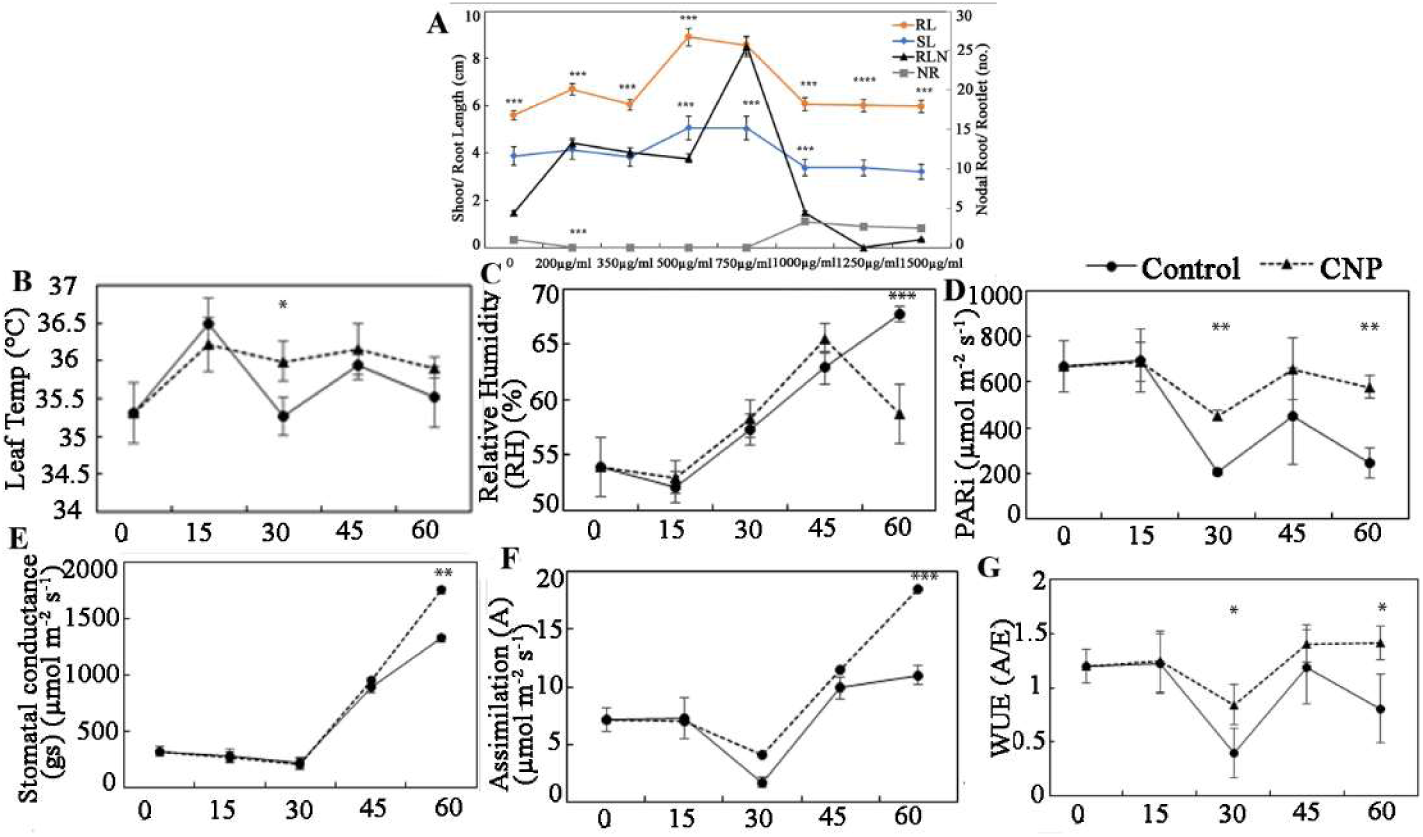
Dosage dependence effect and Leaf gaseous exchange, water relation parameters of CNP exposure. (A): Dosage dependence effect of Carbon nanoparticle exposure on seedling growth. Seedlings were grown on 50 ml Murashige and Skoog (MS) medium containing different concentration of CNP including 200, 350, 500, 750, 1000, 1250 and 1500 μg/mL) for 7 d. On the 8th day, seedling growth parameters such as shoot length (SL), root length (RL), rootlet number (RLN), Nodal roots (NR) were measured. All data presented were from minimum 30 seedlings in each experiment with 2 replications. Significances of values were obtained from one-way ANOVA using the Turkey’s multiple comparison in the Prism version 7.0 software, and were represented as ***, *P* ≤ 0.001. (B-G): Leaf gaseous exchange and water relation parameters in the CNP-treated rice plants. Leaf gaseous exchange, water relation and photosynthetic parameters were recorded in CNP-treated and untreated plants before each of the four CNP treatments on the 45^th^, 60^th^, 75^th^, 90^th^ after sowing (DAS). The last reading was taken 15 days after the last treatment on 90^th^ day (on the 125^th^ day). Thus, the 45^th^, 60^th^, 75^th^, 90^th^, 125^th^ day measurements were noted as 0-day, 15^th^-day, 30^th^ -day, 45^th^ -day and 60^th^ -day of CNP-treatment respectively. The middle part of top second leaf was used for taking readings. Each data point in the graph is a mean of 108 recordings with 6 plants from each either untreated or CNP-treated plants. For each plant 18 recordings were done from 3 leaves in time series of 5 seconds interval, with 6 readings from each leaf. PAR: Photosynthetically actinic radiation, A: Assimilation, GS: stomatal conductance, E: transpiration rate, VPD: vapour pressure deficit, Ci: CO_2_ concentration inside the leaf, WUE: water use efficiency. Significances of values were obtained from one-way ANOVA using the Turkey’s multiple comparison in the Prism version 7.0 software, and were represented as *, *P* ≤ 0.05, **, *P* ≤ 0.01 and ***, *P* ≤ 0.001.

These results indicated that the optimum concentration of CNP on rice seedlings were between 500 and 750 µg/ml. NR showed opposite trend with in increase after at least 750 µg/ml and was higher till 1500 µg/ml of CNP (Figure 1). This further indicated that NR could be a toxicity response induced due to CNP between 1000 - 1500 µg/ml. Germination percentage on 4^th^ day of rice seeds was homogenously 100% in all concentration indicating that CNP didn’t exert any effect on seed germination or growth of seedlings.

### Effect of CNP treatment on physiological parameters in rice plants

Water relation parameters were assessed after four CNP treatments in the treated and untreated plants to investigate the reason of improved growth and yield of CNP-treated plants (Figure 1B-G)). Photosynthesis parameters such as photosynthetically active radiation inside the leaf (PARi) (Figure 1D), carbon assimilation (A) (Figure 1F) showed significantly higher values in the CNP-treated plants. Water relation parameters such as leaf temperature (Tleaf) (Figure 1B), and water use efficiency (WUE) (Figure 1G) were significantly higher (by 0.4°C ± 0.1°C and 75% respectively) in the CNP-treated plants. Relative humidity (RH) (Figure 1C) was significantly less (i.e. 13.3%) in CNP-treated plants. Vapour pressure deficit (VPD) (Figure 2F) and transpiration rate (E) showed negligible changes between the CNP-treated and the control plants (data not shown). Stomatal conductance was also found to be higher in the CNP-treated plants (Figure 1E). CIRASS3 measurements and analysis showed improvements in the water relation traits in the CNP treated plants as compared to their controls.

**Figure 2.**
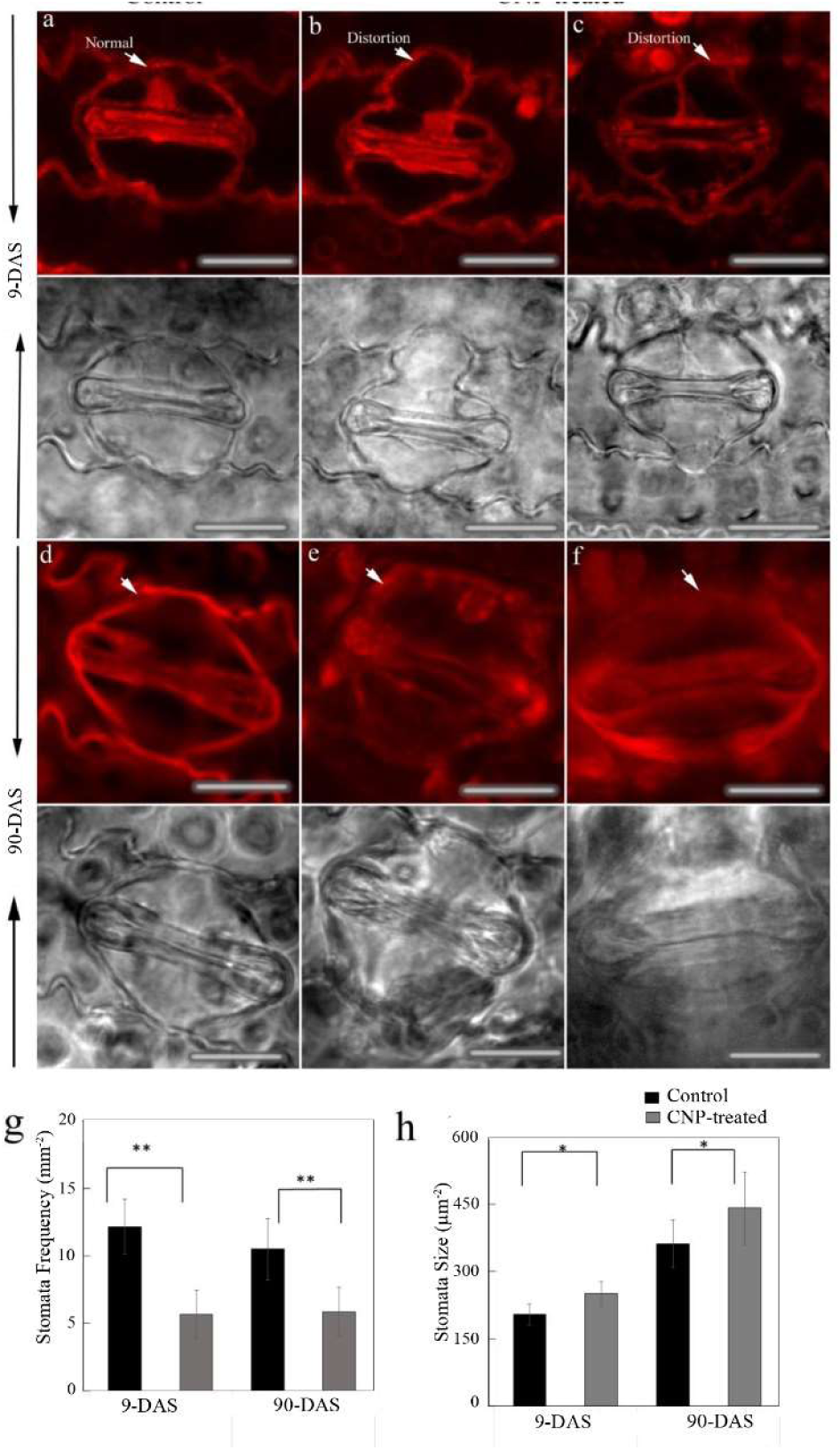
Confocal Microscopic analysis of stomata and cell wall in the CNP-treated rice plants. Leaf Tissue samples from 9d-old seedlings (9 DAS) or 90d-old plants (after 4^th^ CNP treatment) (90 DAS) were fixed with FAA substituted with methanol and stained with propidim iodide to visualize cell wall of stomatal guard cells. Middle portion of the top second leaf was used in case of adult samples. Arrow marks indicate distortions in CNP-treated plants. (A-C): seedlings, (D-F): adult plant leaf. Each representative plant image is placed with its corresponding bright field image below. Scale Bar: 10 µM. G: stomata frequency and H: stomatal size. Size and frequency of stomata were averaged from 6 leaf sections in either control or CNP treated with n≥30.

### Effect of CNP treatment on stomatal parameters in rice plants

As the CNP-treated plants had improved water relation parameters, stomatal analysis was performed using confocal microscopy in the control and CNP-treated samples in the seedlings stage as well as after 4^th^ CNP treatment in plants (Figure 2).

Stomata in the seedling cells showed clear bulging in the cell wall and distortions of the CNP-treated samples which were completely absent in the control samples (Figure 2A-2C). Samples from older plants also showed similar bulging and distortions (Figure 2D-2F). Frequency of stomata per unit area of leaf decreased significantly (∼50%) in both the seedlings and older plant leaves (Figure 2G) with an increase of ∼6% in the size of the stomata (Figure 2H). These results indicated that CNP treatment might have affected to increase stomatal size and decrease in their frequency.

### Analysis of mechanism of CNP uptake

Water uptake by the seedlings with or without CNP in the medium was quantified studying the % increase or decrease of water uptake per seedling (Figure 3).

**Figure 3.**
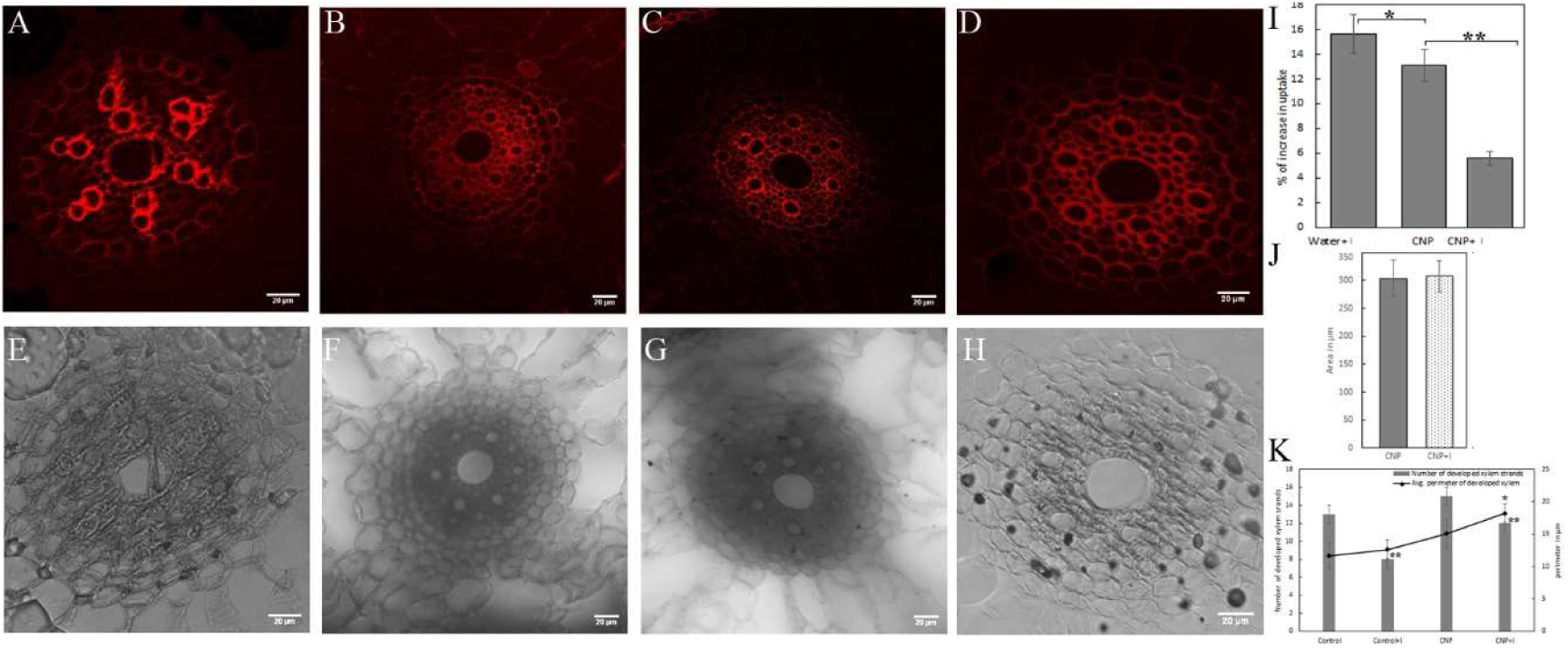
Analysis of CNP-water uptake mechanism in to plant roots. Root tissue samples from 9d-old seedlings with or without CNP treatment were grown in water in air tight plantons. Water uptake/ seedling was calculated as described in materials and methods from water left in the planton and seedling weight. For xylem perimeter and number root samples were fixed, moulded, embedded and sectioned prior to confocal microscopy. CNP aggregates and xylem were analysed from confocal images using Image J software. Data represents mean of 15 focuses at microscope generates from 3 biological replicate root samples in each case. Significances of values were obtained from one-way ANOVA using the Turkey’s multiple comparison in the Prism version 7.0 software, and are represented as *, *P* ≤ 0.05, **, *P* ≤ 0.01.

Addition of CNP in the medium increased water uptake (i.e., 13%) by the seedlings (Figure 3I). To understand if CNP uptake is coupled to water uptake or through exo-/ endo- or pinocytosis, inhibitor nocodazole was added with or without CNP in the medium. Treatment with nocodazole increased the water uptake by 16% than the control with water alone (Figure 3I). Nocodazole treatment in the CNP-treated seedlings reduced the water uptake (6% more than control water alone) as compared to those CNP treated ones (Figure 3I). To investigate the reason of increased water uptake with CNP, number of developed xylem strands and their average perimeter was measured from confocal microscopy images in CNP-treated root samples. Control seedlings with water alone showed 13±1 developed xylem strands with an average perimeter of ∼11.6 µm (Figure 3A, 3E, 3K). Seedlings grown with nocodazole showed reduced number (i.e., 8±1) of developed xylem strands but no change in average perimeter (Figure 3B, 3F, 3K). CNP-treated seedlings showed increase in number of developed xylems (i.e., 14-15 ±1), which were found in 2-3 rings also with increased average perimeter of developed xylem strands of ∼15 µm (Figure 3C, 3G, 3K). Inhibitor application with CNP treatment showed similar number of developed xylem strands as that of CNP, however the average diameter was reduced to ∼10.9 µm indicating reduced water uptake ((Figure 3D, 3H, 3K). Further, an estimate of CNP uptake was made by measuring the area of CNP aggregates in the confocal microscopy images. CNP aggregates in the nocodazole treated samples similar to that with the CNP-treatment alone (Figure 3J).

### Genome-wide expression and pathway analysis of transcripts during CNP-treatment

Our first report in *Arabidopsis thaliana* (*A.thaliana*; At) plants with CNP treatment induced accelerated flowering, photoperiod dependency (Kumar et al., 2018) and in our second study on rice, CNP treatment resulted improved growth and yield along with shade avoidance response (SAR) phenotype and increased plant temperature, albeit with minimal preponement of flowering time (Panigrahy et al., 2021). To understand the basis of the differences in CNP effects in the model dicot and monocot plants, transcriptome analysis and microarray experiment was performed separately from seedling or leaf samples of *A. thaliana* and flag leaf samples of Swarna rice variety respectively (Figure 4).

**Figure 4.**
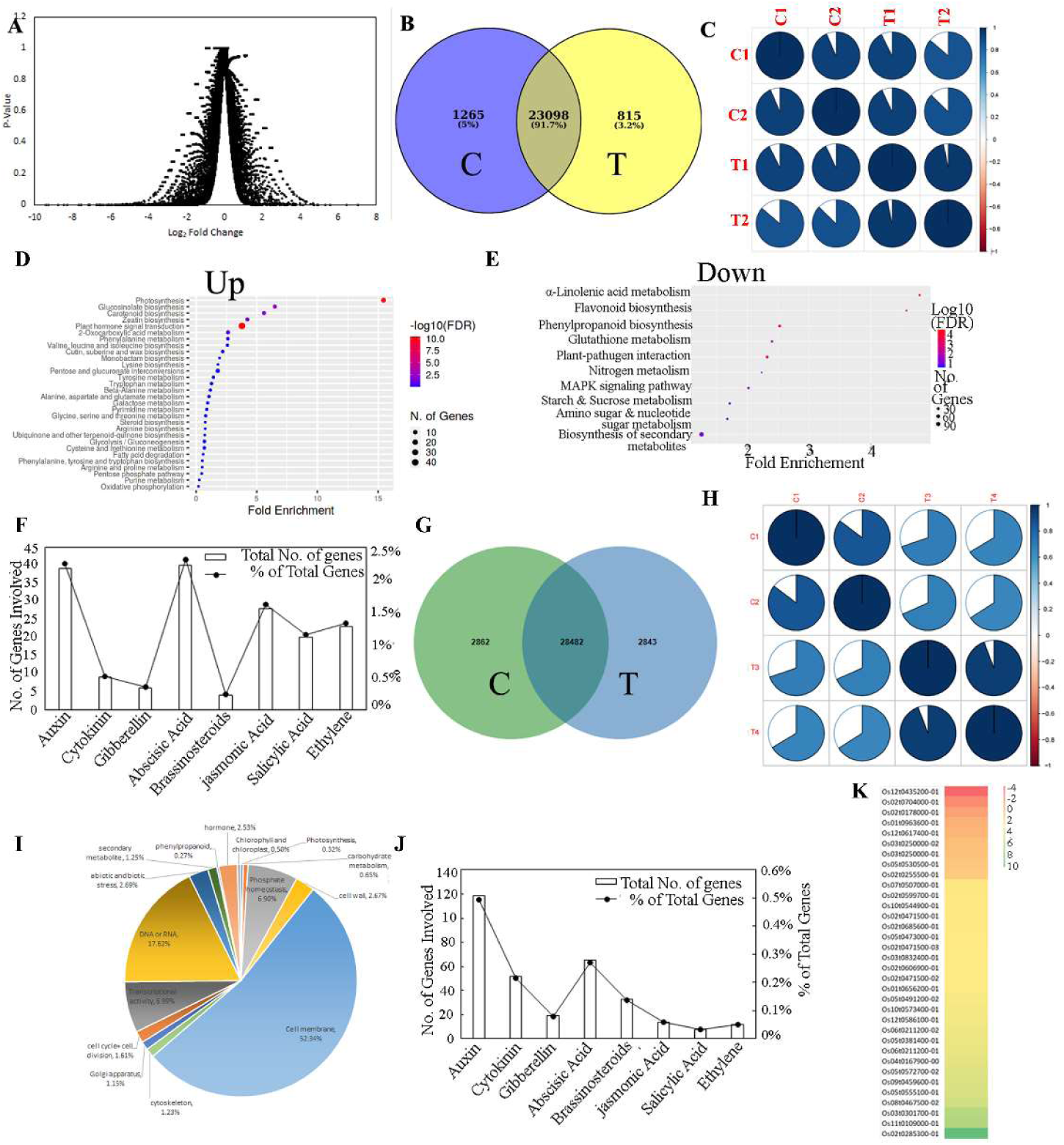
Genome-wide expression analysis in CNP-treated seedlings in rice and *Arabidopsis*. (A-D): Transcriptome sequencing analysis data from *Arabidopsis.* Seedlings were grown on medium without or with 500 µg/ml CNP and sampled on 16^th^ day. (E-H): Microarray gene expression analysis data from rice flag leaf samples on the day of full panicle emergence after 4^th^ CNP treatment. (A): volcano plot of DEG with log_2_ FC cut off 1.0 and P≤0.05, (B): Venn diagram of number of DEG identified, (C-D): Dot plots showing fold enrichment of up and down-DEG, (E): Pathway analysis of eight plant growth hormones in the transcriptome sequencing data (F): Correlation plot among replicates of control and CNP-treatment samples, (G): venn diagram of number of probes identified in microarray experiment, (H): Heat map of 34 selected ABA pathway genes from the microarray data. (I): Pathway analysis of eight plant growth hormones in the microarray DEGs

Each of these experiments were done with 2 biological replicates from control condition or with CNP. At transcriptome volcano plot for transcripts below 1.0 (Figure 4A) showed that highest number of transcripts (37.4%) had p<0.5. Moreover, the replicates of control libraries or the test libraries bear more collinearity (Figure 4C), making them suitable platform for further analysis. Standard analysis of the transcripts (described in Methods section) resulted 23098 differentially expressed transcripts (DEG) which were found in both with and without CNP (Figure. 4B). Pathway enrichment analysis showed the fold-enrichment of the Up- or Down-regulated DEGs (Figure 4D, 4E) with the number of genes of the top 30 pathways. Among the up-regulated DEGs, ‘Plant hormone signal transduction pathways’ was highly enriched with fold enrichment 3.78, enrichment FDR value 3.77413E-11 containing 40 genes in list (Figure 4D). And among the down-DEGs, unsaturated fatty acids such as ‘α-linolenic acid metabolism’ (fold enrichment 4.79, enrichment FDR 1.68024E-05) followed by ‘phenylpropanoid biosynthesis’ with fold enrichment of 2.52 and enrichment FDR of 5.02073E-05 were prominent (Figure 4E). These results indicated that CNP treatment in *A.thaliana* alleviated lipids and secondary metabolites with decreased alpha-linolenic acid like stress antioxidants. Moreover, it indicated an increased involvement of hormone signal transduction. Pathway category analysis of At transcriptome DEGs showed that abscisic acid (ABA) pathway genes were involved to the highest followed by the auxin pathway genes (Figure. 4F). Microarray analysis from CNP-treated and untreated samples of Swarna rice flag leaf samples resulted 52% to 59% of detection of probes in the control and test arrays, with 28,482 probes detected in common (Figure 4G). Collinearity among the control and test replicates above 0.8 (Figure. 4H) created suitable platform for further analysis. Function and pathway analysis showed that major category of transcripts in the CNP-treated samples were from cell membrane, DNA or RNA, transcriptional activity, phosphate homeostasis genes. Hormone pathway genes acquired 2.53% of the total genes involved (Figure 4I). Cohering to the pathway result of *A.thaliana*, pathway analysis of rice samples revealed that ABA pathway genes were highly enriched being the second most enriched after auxin pathway genes (Figure. 4J). Out of the 34 most prominent ABA pathway genes, 29 were found to be up-regulated in CNP-treated samples (Figure. 4K).

### Comparative study of functional categories of differentially expressed genes (DEGs) among rice and *Arabidopsis*

DEGs from the *At* transcriptome and rice (*Os*) microarray were arranged according to their function (GO annotation), pathway involved (KEGG database) and details obtained from TAIR and RAP-annotations for *Arabidopsis* and rice respectively. These relevant categories included ABA pathway, cold stress response, light signaling and circadian clock, photosynthesis and carbohydrate, ROS detoxification, ion and water transport, stomata size, which were used for the comparative analysis among the two model plant systems (Table S1, Table S2). In ABA pathway, 34 genes were found among which majority (24 genes) were up-regulated in rice. Whereas, in *Arabidopsis* only 12 gene were found to be involved in ABA pathway, most of which (∼66%) are down-regulated. Cold stress response category of genes represented different regulation among *At* and *Os*. In rice ∼54% of these genes were up-regulated, whereas in *At* most of the genes in this category were down-regulated. Genes of light signaling, and circadian clock didn’t represent any significant differences in their numbers of up and down DEGs among rice and *At*. In photosynthesis, chlorophyll and carbohydrate category, all the DEGs were down-regulated in rice while ∼63% of DEGs were up-regulated in *At*. DEGs of ROS detoxification category presented contrasting pattern with majority being down-regulated in *At* and up-regulated in rice. However, the transcript levels of superoxide dismutase and catalase in *At* data were upregulated (Table S2). Also, the 11 class III Peroxidases found in rice microarray data were all up-regulated indicating these may act to detoxify the ROS in CNP-treated samples. The DEGs of ion, solute and water transport category were synergistically presented being majority of them down-regulated in both *At* and rice. DEGs of stomata size were up-regulated in both At and rice. These results indicated that CNP treatment induced differential effect on regulation of genes of different functional categories in *At* or rice.

### Effect of CNP treatment on endogenous Abscisic acid content and synthesis

To understand if CNP treatment with abscisic acid (ABA) has any effect on seedling root and shoot growth, (ABA) was added to the growth medium to observe its effect on seedling phenotype. ABA is known to slow down the root and shoot growth in rice (Chen et al., 2006). ABA treatment without or with CNP resulted comparable % of decrease in SL and RL respectively (Figure 5A).

**Figure 5.**
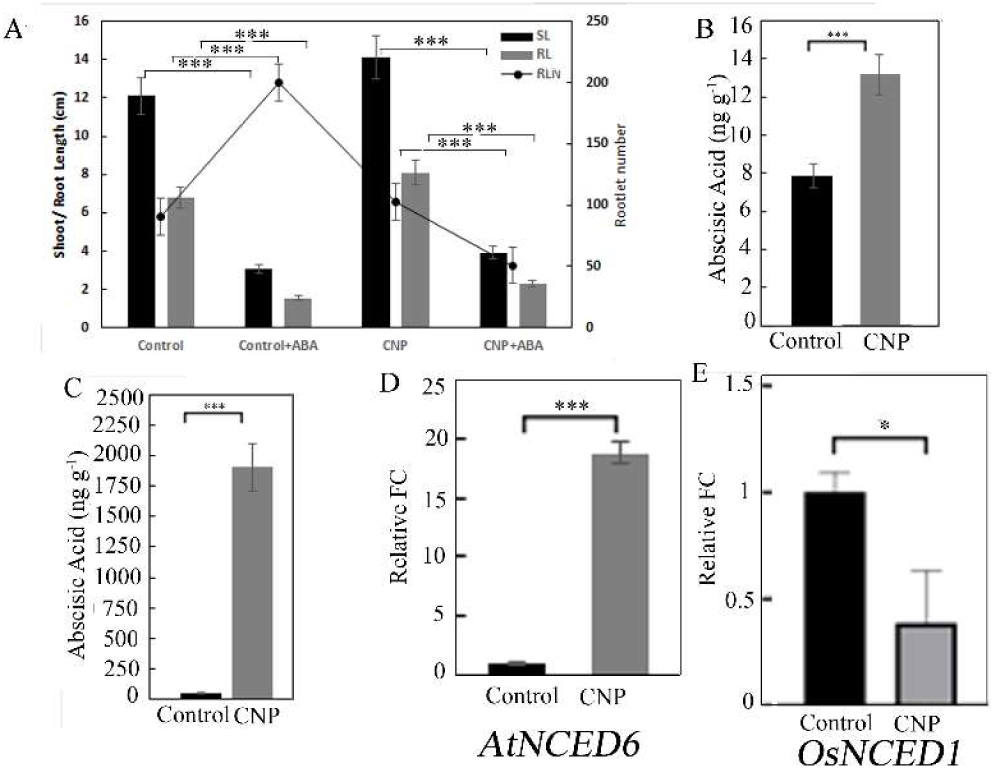
Effect of ABA treatment and endogenous ABA quantification in CNP-treated seedlings. (A) Seedling phenotype was assayed in 14d-old seedlings after 10 µM ABA treatment on the 5^th^ day. (B) Endogenous ABA content was assayed in seedlings and (C) adult leaf samples. Top second leaf was used for lyophilization. (B-C): data represented is a mean of three replicative measurements. Transcript expression levels of ABA biosynthesis gene in (D) Arabidopsis *9-cis-epoxycarotenoid dioxygenase 5* (*AtNCED6*) and (E) rice *9-cis-epoxycarotenoid dioxygenase 1* (*OsNCED1*) were determined using qRT-PCR. Transcript levels of ACTIN was used to normalise transcript levels. Reactions were done in triplicates from two biological replicates. Data represents as the mean ±SEM. The relative quantification of genes was analyzed using standard 2^-ΔΔct^ method. For relative fold-change, the respective expression value for the same gene from untreated plant leaf samples was taken as 1. Statistical significance was obtained from one-way ANOVA using the Turkey’s multiple comparison in the Prism version 7.0 software, and were represented as *, *P* ≤ 0.05, **, *P* ≤ 0.01 and ***, *P* ≤ 0.001.

While SL decreased 74.6% and 72.2%, RL decreased 77.2% and 71.6% in case of control and CNP-treated seedlings respectively (Figure 5A). RLN in the CNP-treated seedlings decreased to nearly half, however surprisingly, the control seedlings showed significant increase of ∼2.2-fold in the RLN with ABA treatment (Figure 5A). These results indicated that CNP induced growth effects could alter endogenous ABA levels. Endogenous ABA in seedlings showed that CNP-treated seedlings had nearly double amount of endogenous ABA than the untreated ones (Figure 5B). Congruently, flag leaf samples of CNP-treated plants were found to have ∼40-fold higher ABA content than the untreated ones (Figure 5C). Additionally, the three other defense hormones such as jasmonic acid (JA), salicylic acid (SA) and oxylipins (oxy) were also found to be higher in the CNP-treated plants (Suppl. Figure 2). Transcript expression analysis of ABA biosynthesis gene *9-cis-epoxycarotenoid dioxygenase 5* (*AtNCED6*) in *Arabidopsis* seedlings (Figure 5D) and *9-cis-epoxycarotenoid dioxygenase 1* (*OsNCED1*) in rice flag leaves (Figure 5E) were validated to support the above observation of ABA content. *AtNCED6* relative expression level was increased to ∼18-fold and *OsNCED1* relative expression level was decreased to ∼0.39-fold after CNP treatment.

### Transcript expression analysis to understand CNP-treated plants’ phenotype

To have insights into CNP-treated plant’s phenotype, transcript expression of different selected genes from photosynthesis, calvin cycle, cold tolerance, stomata size and frequency, genes for ABA pathway were studied using qPCR in CNP-treated plants in comparison with their control plants. The concordance of the transcript expression levels of majority of the genes analysed from the rice microarray with the qRT-PCR fold change provided evidence of sound genomic data set (Suppl. Figure 3A). Photosynthesis related gene *OXYGEN EVOLVING COMPLEX1* (*OsOEE1*) was significantly down-regulated to 0.25-fold in CNP-treated samples compared to the controls (Figure 6A).

**Figure 6.**
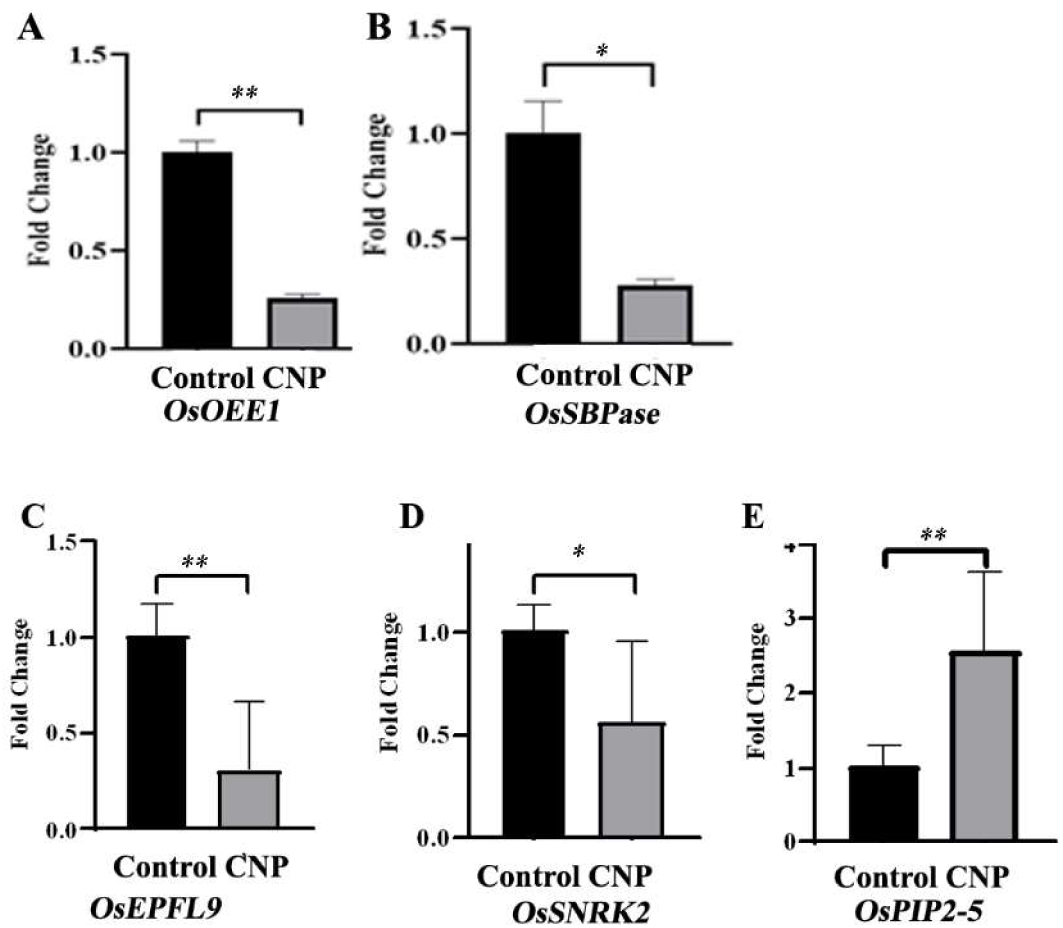
Transcript expression analysis in CNP-treated rice plant samples. Transcript expression levels of genes involved in photosynthesis, cold tolerance, stomata and water relations were studied using qRT-PCR in flag leaf samples after 4^th^ CNP treatment. (A): *OXYGEN EVOLVING COMPLEX1* (*OsOEE1*), (B): *SEDOHEPTULOSE 1-7-BI-PHOSPHATE* (OsSBPase), (C): *EPIDERMAL PATTERNING FACTOR-LIKE 9* (*OsEPFL9*), (D): *SUCROSE NON-FERMENTING-1 RELATED PROTEIN KINASE 2* (*OsSNRK2*), (E) *PLASMA MEMBRANE INTRINSIC PROTEIN 2-5* (*OsPIP2-5*). Transcript levels of ACTIN was used to normalise transcript levels. Reactions were done in triplicates from two biological replicates. Data represents as the mean ±SEM. The relative quantification of genes was analyzed using standard 2^-ΔΔct^ method. For relative fold-change, the respective expression value for the same gene from untreated plant leaf samples was taken as 1. Statistical significance was obtained from one-way ANOVA using the Turkey’s multiple comparison in the Prism version 7.0 software, and were represented as *, *P* ≤ 0.05, **, *P* ≤ 0.01 and ***, *P* ≤ 0.001.

*SEDOHEPTULOSE 1-7-BI-PHOSPHATE* (OsSBPase) (Suzuki et al., 2021, Feng et al., 2007) was significantly down-regulated to 0.27-folds in CNP-treated samples compared to their controls (Figure 6B). These results indicated that the input to the calvin cycle through the *OsSBPase* as well as the photosynthetic yield, as indicated through *OsOEE1* (Heide et al 2004) transcript expression levels, were deteriorated in the CNP-treated plants compared to the untreated plants. To confirm the microscopic observations of stomatal parameters in the CNP-treated plants, genes related to stomatal size and frequency including *EPIDERMAL PATTERNING FACTOR-LIKE 9* (*OsEPFL9*) and *SUCROSE NON-FERMENTING-1 RELATED PROTEIN KINASE 2* (*OsSNRK2*), were studied for their transcript expression levels. *OsEPFL9* transcript expression level was found to be reduced to ∼0.31-folds in the CNP-treated samples compared to the controls (Figure 6C). *OsSNRK2* transcript expression level in the CNP-treated plants was found to be reduced to nearly half (∼0.55-fold) compared to their controls (Figure 6D). To understand the molecular basis of the altered water relation parameters in the CNP-treated plants, water use efficiency (WUE) related aquaporin gene *PLASMA MEMBRANE INTRINSIC PROTEIN 2-5* (*OsPIP2-5*) transcript level was studied comparatively. *OsPIP2-5* transcript level was significantly increased to ∼2.5-fold in the CNP-treated seedlings compared to the controls (Figure 6E). To confirm the involvement of CNP in abiotic stress response *PROTEIN PHOSPHATASE 51* (*OsPP51*) transcript level was studied in CNP-treated seedlings compared to their controls. *OsPP51* level was up-regulated to ∼5-folds in the CNP-treated samples (Table S1) indicating improved abiotic stress compared to the controls.

### Effect of different abiotic stresses in CNP-treated seedlings

Improved water relation parameters, bigger sized stomata and altered endogenous ABA content urged to study the effect of different abiotic stresses in CNP-treated seedlings. To this end, various stresses (i.e., cold at 18°C, high temperature at 35 °C, salinity stress and induced drought using PEG) were imposed in separate experiments on seedlings with or without CNP in growth medium.

Changes in the RL under all these stresses were insignificant, hence, effects were studied in SL and RLN only. Seedlings showed 16.5% increase in SL and ∼47.1% increase in RLN due to CNP treatment (Figure. 7A). This data served as the control for the abiotic stress treatments tested. Cold treatment for 7 d showed significant decrease in SL and RLN (Figure. 7A). While SL decrease ∼64% without CNP, it decreased ∼60% with CNP due to cold stress. RLN also showed higher decrease (∼98%) without CNP and ∼91.8% decrease with CNP due to cold temperature. These results indicated that CNP treatment could partly rescue the drastic effects on seedling growth during cold stress. PEG treatment creates osmotically induced drought environment at cellular level (Purbajanti et al., 2019). PEG treatment showed marginal insignificant increase in SL in both untreated and CNP-treated seedlings (Figure. 7A). Effect of PEG treatment showed significant contrasting response on RLN with or without CNP, i.e., 12.7% decrease without CNP and 9.2% increase with CNP treatment as compared with the control and CNP seedlings respectively. These results indicated that CNP treatment effectively counteracts the PEG induced cellular drought to show increase in RLN. Salt stress induced decrease in SL, RL and RLN both with and without CNP (data not shown). The % decrease due to salt stress was comparable, insignificant and within the range. High temperature treatment resulted decrease in SL, RL and increase in RLN (data not shown), however the differences were insignificant, therefore were not of interest. Despite germination, growth of seedlings was hampered under low temperature from 6^th^d-14^th^d. But, CNP-treated seedlings SL showed less effect as compared to the untreated ones (Figure.7B). These results indicated that CNP treatment has selective effects to countereffect the stress caused due to cold temperature or PEG treatment.

**Figure 7.**
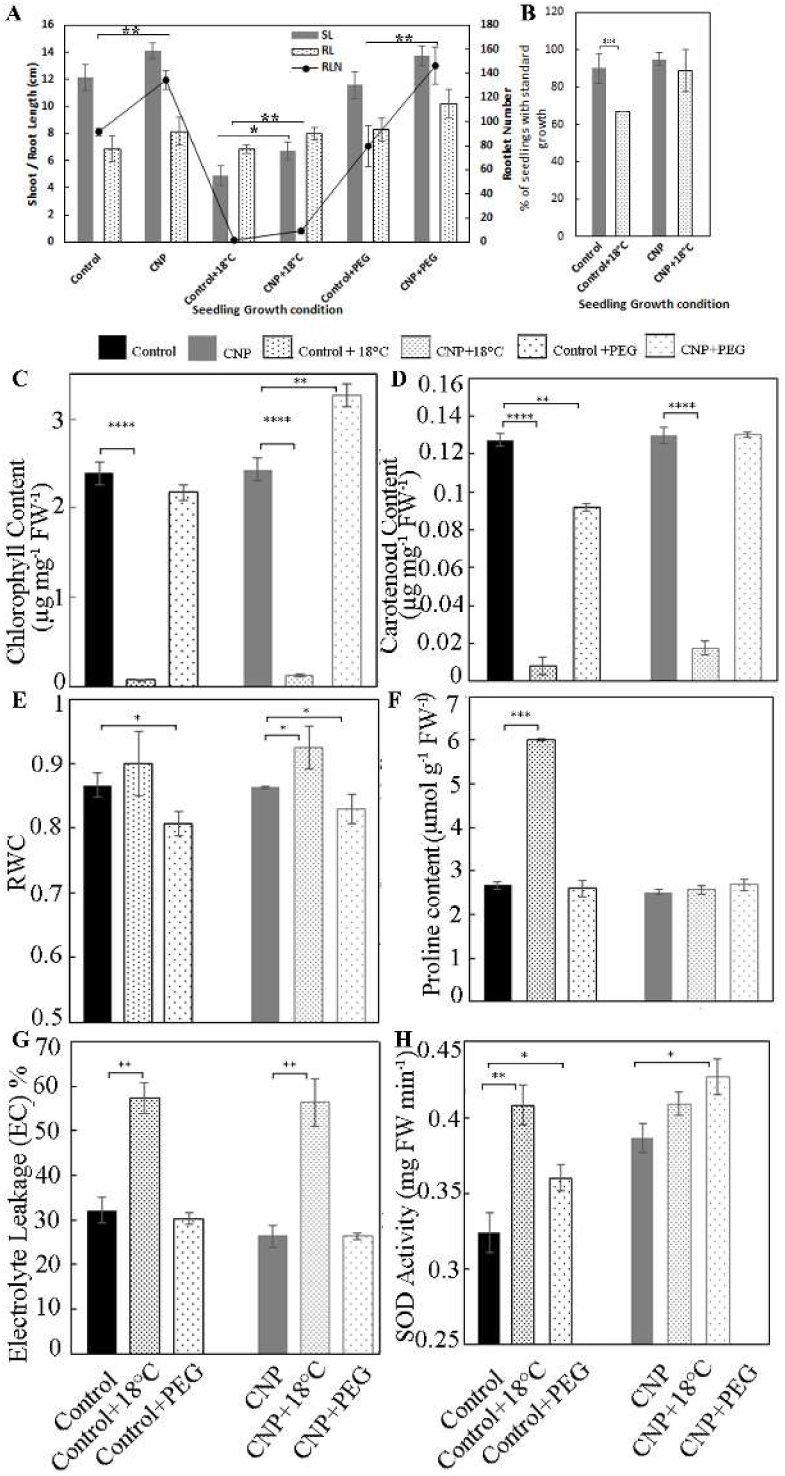
Effect of different abiotic stresses in CNP-treated seedlings. (A-B): Five-day-old seedlings were carefully transferred on to medium containing PEG or salt and grown for 7 more days. On 13^th^ – 14^th^ day, seedling growth parameters such as SL, RL, RLN were phenotyped. For cold treatment, seedlings were grown one more day on MS medium before treatment for attaining higher growth to compensate the growth retardation due to cold. All data presented were from minimum 30 seedlings in each experiment with 2 replications. (C-H): Physiological and Biochemical characteristics of Cold and PEG stress response in CNP-treated seedlings. Physiological and biochemical characteristics of seedlings with cold stress (18°C) or artificial drought due to PEG treatment for 7 days were assayed in 13d-old seedlings (14-day old in case of cold treatment). Seedlings were grown on MS or CNP medium till initial 4d before stress treatment. C: Chlorophyll content, D: carotenoid content, E: RWC content, F: proline content, G: Electrolyte leakage %, H: Superoxide dismutase activity. These experiments were done thrice from leaves grown in at least 3 biological replicates. Data represented is a mean of 9 readings with 3 replications each time. Statistical significance was obtained from one-way ANOVA using the Turkey’s multiple comparison in the Prism version 7.0 software, and were represented as *, *P* ≤ 0.05, **, *P* ≤ 0.01 and ***, *P* ≤ 0.001.

### Countereffect of CNP-treatment on cold- and PEG-treatment induced stress in seedlings

To ascertain the seedlings growth effects observed under cold-stress and PEG treatment and the countereffects of CNP on it, various physiological and biochemical parameters were studied in 13 d-old (14 d-old in case of cold stress) seedlings with or without CNP (Figure 7C-H). Cold stress resulted drastic decrease in chlorophyll content (∼32.8-fold and ∼18.2-fold in case of control and CNP-treated seedlings respectively), with higher amount of chl left in CNP-treated seedlings (Figure 7C). Similarly, carotenoid content was also drastically reduced due to cold stress with ∼16.1-fold and ∼7.3-fold in case of control and CNP-treated seedlings respectively and lead to similar pattern of higher carotenoid content in CNP-treated seedlings left after cold stress (Figure 7D). Relative water content (RWC) was not significantly different in the control seedlings with or without cold stress, however it was 7.1% increased (p ≤ 0.05) in the CNP-treated seedlings with cold stress when compared to without cold stress (Figure 7E). Electrolyte leakage (EC) % was significantly higher in the control as well as the CNP-treated seedlings after cold stress (Figure 7G) (1.78-fold and 2.1-fold respectively). Proline accumulation showed drastically different pattern among the CNP-treated and control seedlings (Figure 7F). In the control seedlings, proline content was ∼2.25-fold higher after cold stress as compared to the respective unstressed ones, whereas it showed nearly no change in their values in CNP-treated seedlings due to cold stress. Reactive oxygen species scavenging enzyme superoxide dismutase (SOD) activity was significantly higher in control (i.e., 5.7%), however in the CNP-treated seedlings SOD activity was not much increased after cold stress (Figure 7H). These results indicated that the effect of cold stress was alleviated due to CNP treatment seedlings with higher chlorophyll, carotenoid, RWC, lesser SOD activity and nearly no more proline accumulation as compared to control seedlings.

PEG treatment for 6 d did not show significant decrease in chlorophyll in the control seedlings, whereas it showed ∼ 34.1% increase (p ≤ 0.01) in chlorophyll in CNP-treated seedlings (Figure 7C). Carotenoid content showed significant decrease by 27.7% in the control seedlings due to PEG treatment, whereas it showed nearly no change in the CNP-treated seedlings (Figure 7D). RWC decreased marginally but significantly in both control and CNP-treated seedlings (6.8% and 3.8% respectively) after PEG treatment (Figure 7E). Proline accumulation (Figure 7F) and EC% (Figure 7G) were insignificant in both control and CNP-treated seedlings after PEG treatment. Superoxide dismutase (SOD) enzyme activity was higher after PEG treatment in both control and CNP-treated seedlings, however the CNP-treated seedlings it was higher than the control ones (Figure 7H). These results indicated that after PEG treatment, CNP-treated seedlings retained drought resistance characteristics but only in some parameters including chlorophyll, carotenoid content, RWC.

### Assessment of cold and drought stress tolerance in rice plants in reproductive stage

To confirm the cold and PEG-induced-drought tolerance results obtained in the seedlings with CNP treatment, cold and drought stress was imposed in rice plants in the reproductive stage (after 4^th^ CNP treatment).

Cold stress at 10 °C for 10 d affected the cut leaves of untreated and CNP-treated leaves in a similar manner (Suppl Figure 4). Yellowing of leaves, cold-induced lesions and drying progressed in similar fashion in untreated and CNP-treated leaves (Suppl Figure 4A, 4B). Also, a recovery period of 24 h at room temperature was able to regain the greenness of leaves in untreated and CNP-treated leaves in a similar manner (Suppl Figure 4C). However, chlorophyll and carotenoid content in the cold-treated leaves were higher in the CNP-treated leaves than the untreated ones (Suppl Figure 4D-E). Further parameters were not analyzed as the leaves of untreated and CNP-treated leaves were apparently similar after cold stress.

Drought tolerance was tested by withholding water in control and CNP-treated pots (Figure 8A). Water-stress (WS) for 36 h showed clearly resistant plant phenotype in the CNP-treated plants with greener stem, expanded and greener leaves than the control plants (Figure 8B). After 48 h of water stress (Figure. 8C) and 24 h after re-watering, the CNP-treated plants had clearly more greener leaves (Figure 8D). Chlorophyll content showed ∼41% and ∼61.2% decrease in both the untreated and CNP-treated plants respectively after WS. However, CNP-treated leaves retained the higher chlorophyll content after WS (Figure 8E). Carotenoid content showed to be increased ∼10 times after WS in both control and CNP-treated leaves with the later having the higher values after WS. RWC decreased drastically to nearly half of its starting value in the control plants, however it retained 82% in CNP-treated plants after WS (Figure 8F). MDA content sharply increased by ∼ 4.2-fold of its initial value in the control plants, however MDA content showed only ∼2.2-fold increase in CNP-treated plants after WS (Figure 8G), thus showing less MDA accumulation in CNP-treated plants than the control ones after WS. Proline accumulation showed opposite pattern in control or CNP-treated plants after stress. While proline content increased to ∼1.5-fold in the control plants, it decreased non significantly in CNP-treated plants after WS (Figure 8H).

**Figure 8.**
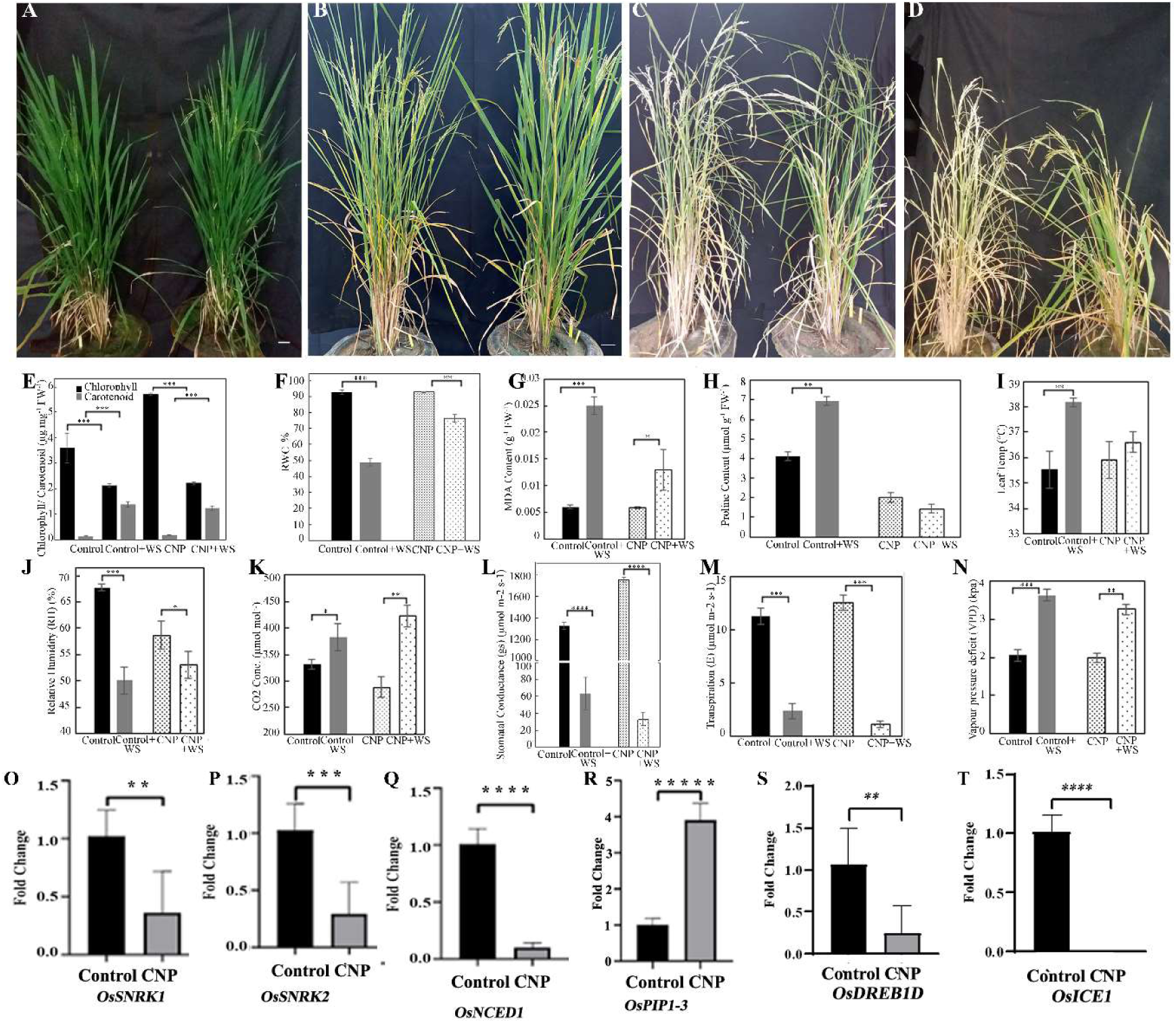
Effect of water stress in CNP-treated plants. (A-N): Water was withheld for 48 h after4^th^ CNP treatment and was compared with the untreated control plants. (A): Plants before water stress, Left plant is control and right one is CNP-treated, (B)and (C): plants after 36- and 48-hours of water withheld respectively, (D): plants after 24-hours of recovery from water stress. Physiological analyses such as (E) chlorophyll and carotenoid content, (F) relative water content, (G) malondialdehyde content, (H) proline content, (I) leaf temperature, (J) relative humidity, (K) leaf internal CO2 concentration, (L) stomatal conductance, (M) transpiration rate and (N)vapor pressure deficit. were done from water-stresses leaf (top 2^nd^) samples. Data was obtained from 3 replicates each from 6 individual plants of untreated or CNP-treated plants. (O-T): Transcript expression analysis in CNP-treated rice plant samples under cold or water stress. Transcript expression levels of genes involved in ABA biosynthesis, ABA-regulated stomatal closure and cold tolerance, stomata and water relations were studied using qRT-PCR in flag leaf samples after 48-hours of water stress. (O): *SUCROSE NON-FERMENTING-1 RELATED PROTEIN KINASE 1* (*OsSNRK1*), (P): *SUCROSE NON-FERMENTING-1 RELATED PROTEIN KINASE 2* (*OsSNRK2*), (Q): *9-cis-EPOXYCAROTENOID DIOXYGENASE 1* (*OsNCED1*) (R) *PLASMA MEMBRANE INTRINSIC PROTEIN 1-3* (*OsPIP1-3*), (S) *DEHYDRATION RESPONSIVE ELEMENT BINDING PROTEIN 1D* (*OsDREB1D),* (T): *INDUCER OF CBF EXPRESSION1* (*OsICE1)*. Transcript levels of ACTIN was used to normalise transcript levels. Reactions were done in triplicates from two biological replicates. Data represents as the mean ±SEM. The relative quantification of genes was analyzed using standard 2^-ΔΔct^ method. For relative fold-change, the respective expression value for the same gene from untreated plant leaf samples was taken as 1. Statistical significance was obtained from one-way ANOVA using the Turkey’s multiple comparison in the Prism version 7.0 software, and were represented as *, *P* ≤ 0.05, **, *P* ≤ 0.01 and ***, *P* ≤ 0.001.

Water relation and related parameters also investigated in the CNP-treated plants in comparison with control plants after WS. Leaf temperature increased by ∼2.5°C in control while insignificantly in CNP-treated plants after WS (Figure 8I). Relative humidity (RH) dropped ∼37% in the control plants while it decreased insignificantly in CNP-treated plants after WS (Figure 8J). Leaf CO_2_ concentration increased only ∼15% in control plants, whereas in CNP-treated plants it increased to ∼1.4 fold, maintaining the highest CO_2_ concentration after WS (Figure 8K). Stomatal conductance was higher in CNP-treated plants in well-watered conditions. WS lead to drastic reduction of stomatal conductance in control plants up to ∼20-fold. However, CNP-treated plants showed even higher (∼53-fold) reduction of stomatal conductance under WS conditions (Figure 8L). Transpiration rate presented similar scenario like stomatal conductance showing drastic reduction after WS (Figure 8M). However, the CNP-treated and untreated plants had insignificant differences in their transpiration rate values both before and after WS. Vapor pressure deficit (VPD) was comparable between control and CNP-treated plants under well-watered conditions, however, WS increased the VPD to nearly two-fold with insignificant differences between control and CNP-treated plants after WS (Figure 8N). These results indicated that CNP treatment in the rice plants could help to better withstand water stress by maintaining high chlorophyll carotenoid content, lower MDA, proline content, lower leaf temperature and lower stomatal conductance, transpiration rate.

### Transcript expression analysis after water and cold stress in CNP-treated plants

To confirm the drought and cold stress tolerance phenotype equipped with biochemical and physiological characteristics in the CNP-treated plants during above stresses, transcript expressions of some ABA signaling, stomatal parameters and aquaporin genes were studied from flag leaf samples of CNP-treated plants after cold or water stress in comparison with their controls under well-watered condition (Figure 8O-8T). Expression levels of ABA signaling components which are involved in stomatal closure such as *OsSNRK1* and *OsSNRK2* were down to ∼0.36 and ∼0.29-folds respectively after water stress in the CNP-treated samples as compared to their controls (Figure 8O, 8P). ABA biosynthesis gene *OsNCED1* expression was also down-regulated significantly by 10-fold in the CNP-treated samples (Figure 8Q). Further, aquaporin gene *OsPIP1-3* expression was found to be significantly up-regulated to ∼3.9-folds in the CNP-treated samples as compared to their controls (Figure 8R). These results indicated that CNP-treatment could alter ABA pathway, stomatal and water transport parameters to withstand water stress. To validate the cold stress tolerance parameters with CNP treatment observed at the seedlings stage and to access the cold tolerance at molecular level, transcript expression of *INDUCER OF CBF EXPRESSION1* (*OsICE1) (BHLH116)* and *DEHYDRATION RESPONSIVE ELEMENT BINDING PROTEIN 1D* (*OsDREB1D)* were studied in rice flag leaf samples from CNP-treated and untreated plants after cold stress. *OsICE1* expression was found to be significantly (p<0.0005) down-regulated to nearly zero in the CNP-treated samples (Figure 8S). *OsDREB1D* was also significantly (p<0.01) down-regulated in the CNP-treated samples, however the down-regulation was 0.25-folds as that of the controls (Figure 8T). These results confirmed that cold tolerances response genes were down-regulated in the CNP-treated leaf samples.

## Discussion

External exposure of carbon nanoparticle in the growth medium or pot soil of *Arabidopsis* (Kumar et al., 2108) and rice (Panigrahy et al., 2021) respectively resulted some synergistic while some differential responses in our two previous reports. In *Arabidopsis* it acted as a flowering time accelerator with photoperiod dependency, while in rice it resulted improvement in growth and yield. The synergistic effects included involvement of red-light photoreceptor Phytochrome B with shade avoidance like scenario in both the model plants. The present study not only finds the causes of differential responses due to CNP to some extent in *At* and rice, but also it defines the specific constraints of its use in rice plants and highlights its agronomic beneficial effects with respect to abiotic stress tolerance in rice. For this purpose, several physiological, biochemical, microscopic analyses including genomic analyses were accessed in both the model plant systems. CNP concentrations at 500 µg/mL -750 µg/mL were found to be the optimum dosages showing best seedling growth among the 7 different concentrations tested in the range between 200 µg/mL – 1500 µg/mL with rootlet number being the best parameter to study the seedling response. Results of CNP and water uptake revealed that perimeter of xylem had an important role in water uptake. Endocytosis inhibitor could not affect water uptake in the control seedlings. However, it could reduce the water uptake when CNP was present the medium. Thus, CNP-coupled to water uptake was endocytosis mediated. However, CNP aggregate was not affected with or without nocodazole. CNP alone could be taken up using pinocytosis or energy consuming processes which requires further study.

CNP exposure induced specific resistance to cold stress tolerance at seedling stage with most of the physiological and biochemical parameters according to cold tolerant response. However, in mature rice plant leaves cold tolerance as not very prominent in rice leaf assay. Thought nearly half of the genes in rice microarray were up-regulated, most of the transcripts cold stress response pathway were down regulated in *Arabidopsis*. Cold tolerance response was exhibited in rice seedlings with CNP-treatment as verified through their SL and biochemical parameters including Chl, carotenoid MDA and EC. However, the yellowing and cold-induced lesions in the flag leaves due to cold treatment progressed in a similar manner in the CNP-treated as well as untreated samples. The higher Chl content in the cold-stressed leaves could be due to the chl degradation metabolites. Overexpression of rice *DEHYDRATION RESPONSIVE ELEMENT BINDING 1D* (*OsDREB1D*) or *INDUCER OF CBF EXPRESSION 1* (*OsICE1*) showed cold stress tolerance (Zhang et al., 2009; Dian-jun et al., 2008) in previous reports. Transcript expression of cold pathway genes (*OsICE1, OsDREB1D*) in our study showed down-regulation, indicated that albeit the cold stress tolerance that was exhibited in the seedlings stage, CNP-induced cold stress tolerance was not efficient enough to be exhibited in reproductive stage. Our previous report of CNP treatment in rice resulted a clue of involvement of jasmonic acid and auxin pathway genes for CNP response (Panigrahy et al., 2021). This indication of JA and auxin involvement was proved in the present study by the highly elevated JA content in rice leaf samples (Suppl Figure 2) and also by the major involvement of the auxin pathway gene in the genomic data of *At* and rice. The discrepancies in the CNP responses among the *At* and rice could be due to the differential involvement of ABA pathway, photosynthesis and carbon metabolism pathway genes among the two model plants. Nevertheless, almost all the Protein phosphatase 2C (PP2C) found in the microarray data were up-regulated could clearly indicate strong positive involvement of ABA pathway for CNP phenotype in rice. <u>S</u>edoheptulose-1,7-<u>B</u>isphosphat<u>ase</u> (OsSBPase) is known as an important rate limiting enzyme for fuelling carbon input to the calvin cycle (Feng et al., 2007) and the <u>O</u>xygen <u>E</u>volving <u>E</u>nhancer protein 1 (OEE1) of photosystem II is known to control the photosynthesis rate by being involved in ROS scavenging and repair of PSII subunits (Heide et al., 2004). Presently, the decreased expression levels of these photosynthesis genes indicated that the input to the calvin cycle (i.e. SBPase) as well as the yield of the light reaction (OEE1) was less in CNP-treated plants despite higher carbon assimilation rate (Figure 2). However, the CNP-treated plants exhibited an increase in yield/plant as shown previously. This could be due their higher expression of deposition in the grain, higher rate of grain filling as shown before (Panigrahy et al., 2021). These observations were supported by the elevated expression of *SOLUBLE STARCH SYNTHASE IIA* (*OsSSIIa*) and *SOLUBLE STARCH SYNTHASE IIIA* (*OsSSIIIa*) (Table S1). Compensation to the photosynthetic calvin cycle could be provided by the RuBisCO subunit binding-protein (*OsRBCS1*), which was highly over-expressed in the CNP-treated plants (Table S1). Congruently, *Arabidopsis* photosynthesis and carbohydrate pathway showed majority of the genes up-regulated (Table S2). These results explained the higher yields in the CNP-treated plants.

CNP-treated plants were found to have increased ABA content supported by increased expressions of ABA biosynthesis gene in *Arabidopsis* (*AtNCED6*) and decreased expression (*OsNCED1*) in rice. *OsNCED1* expression was shown to be down-regulated during water-stress and in turn stimulate ABA accumulation in rice through a probable feedback inhibition (Changan et al., 2018). *Arabidopsis AtNCED6* expression was shown to increase during water stress (Seiler et al., 2014). Hence, our results of increased ABA content were due to increased ABA biosynthesis in *Arabidopsis* and feedback regulation due to *OsNCED1* in rice. Pyrabactin resistance 1 like (PYL) genes are known to be the ABA receptor. Out of the 3 *PYL* genes found in rice microarray data, 2 of them were up-regulated mildly in microarray data. Despite the two *PYL* genes found in *Arabidopsis* and one rice PYL genes being down-regulated, the positive involvement of ABA pathway could be suggested due to the existence of nearly 13 *PYL* genes in rice (Miao et al., 2018) and 14 *PYL* genes in *Arabidopsis* (Gonzalez-Guzman et al., 2012), which are presently not detected our experiment. Promoting root hair growth is characteristic effect of ABA in rice (Chen et al., 2006). CNP-treated seedlings had higher number of rootlets and CNP-treated adult plants had larger root volume (Panigrahy et al., 2021), it indicated that these effects could be due to elevated ABA content due to CNP. To support this hypothesis, we observed no rootlets in CNP-treated seedlings while having increase in RLN in the untreated seedlings with ABA treatment. It could be further inferenced that higher ABA content in CNP-treated plants could result improved water relation parameters including higher stomata size, stomatal conductance and WUE. Sucrose non-fermenting 1-related kinase 2 (OsSNRK2) is a key mediator regulating plant abiotic stress and is involved in stomata closure (Kulik et al., 2011; Dey et al., 2016). OsSNRK2 is known to be a negative regulator of ABA signaling. Epidermal patterning factor-like 9 (OsEPFL9) is known to be involved in stomatal development and frequency (Yin et al., 2017). Protein phosphatase OsPP51 is known to modulate abiotic stress response in rice (Singh et al., 2010). Aquaporins are the major determinants of water use efficiency in rice (Guo et al., 2006). Our observations of improved water relation parameters were also backed by the increased expression of stomata frequency gene *OSEPFL9*, *OsPP51*and aquaporin gene *OsPIP2-5* and decreased expression of *OsSNRK2*. Hence, there are stronger evidence of improved water use efficiency in the CNP-treated plants. The CNP-treated plants had better tolerance to water stress as evidenced by their increased RH, VPD and RWC compared to the controls. CNP-treated plants could better withstand water stress also had faster recovery than untreated plants, as evidenced not only from their phenotype but also from the several physiological parameters including higher chlorophyll, carotenoids, proline content. Despite higher MDA content, CNP-treated plants had advantage due to their higher water withholding capacity as evidenced by higher VPD, RH and reduced stomatal conductance and reduced transpiration. These observations were also supported by the increased expression level of *OsPIP1-3* responsible for water balance during drought (Verma et al., 2020). During water stress, ABA signaling mediator controlling stomatal closure *OsSNRK1, OsSNRK2* and ABA biosynthesis gene *OsNCED1* expressions were down indicating up-regulation of ABA pathway which may boost the water stress tolerance of the CNP-treated plants.

There was an interference of CNP with the internal CO_2_ concentration in CNP-treated plants which was observed as reduced Ci in well-watered and increased Ci during water-stress as compared to their controls. It was shown in previous report that carbon nanomaterials in plants can interfere with availability of carbon pool to the photosynthesis. SWNT have been shown to add to the light capture and thereby add to the electron transfer and increase photosynthesis rate (Khatri et al., 2018; Giraldo et al., 2014). Though the CNP could not add to the photosynthetic carbon input in our study, somewhat similar phenomenon may interplay in CNP-treated plants to have the present set of observations of improved WUE. Similar findings of carbon nanomaterials in ameliorating plant water status by adjustment of osmolytes, reduced transpiration by stomatal closure due to abscisic acid signaling with reduced photosynthesis have been discussed (Maswada et al., 2020).

In the present scenario, CNP, due to their black color, helped to hike the plant temperature in the rice plants by absorbing more heat energy. Due to slightly warmer temperature, down-regulation of *PHYB*, the recently known thermo-sensor, created a SAR-like syndrome of growth (Panigrahy et al., 2021). PhyB is known to regulate stomatal density, transpiration rate and drought tolerance negatively in rice, as *OsphyB* mutants exhibited reduced stomatal density with drought tolerance (Liu et al., 2011). The decreased *PhyB* expression levels in our case may help to decrease the stomata density and increase their size by up-regulating the *OsEPFL9* expression levels. PhyB was also shown to maintain the low ABA levels in well-watered as well as high ABA levels under water stress, there by regulating the stomatal closure and stomatal conductance (Gonzalez et al., 2012). In Arabidopsis *At*p*hyB* mutants exhibited increased stomatal conductance thereby less tolerant to water stress than the wild type counterparts. Here it is worth mentioning that *phyB* mutants in rice or *Arabidopsis* resulted contrasting responses to drought, and hence, further investigations are required to deduce conclusions of this contrasting response to drought. With reduced *PHYB* levels in our case, the CNP-treated plants could show drought tolerance due to reduced stomatal conductance, transpiration, lowering leaf temperature, maintaining VPD and RH. This could be explained due to the property of carbon nanoparticles, by which they can act as hydroscopic binders of the intracellular water and help plant cells to hold extra water in different compartments of cell (Ahmad et al., 2020). With enlarged stomata, expression of aquaporin *OsPIP2-5* may be altered and may lead to increase the water uptake. These changes may bring significant improvement with reduced transpiration and stomatal conductance to exhibit water stress tolerance. Further investigations to demonstrate the hydroscopic abilities of CNP in our case can unravel detailed molecular mechanisms underlaying the phenomenon of water stress tolerance in CNP-treated plants. These results of this study is definitely a robust increment in building the fundamentals of nano agriculture, the concept and applicability can be exploited in rice and other crops.

## Materials and methods

### Seedling and Plant growth condition

#### Seedling growth condition

Rice variety Swarna (MTU7029) was used for all experiments in this study. Swarna MTU7029 variety was developed by All India Coordinated Rice Improvement Project (AICRIP) centre, Andhra Pradesh Rice Research Institute (APRRI) and Regional Agricultural Research Station (RARS) Maruteru, Acharya NG Ranga Agricultural University (ANGRAU), Andhra Pradesh, India in 1982 under the IET 5656 (IIRR, 2015; MARUTERU). Seeds were surface-sterilized, stratified at 4 °C for 2 d in dark and sown on 50 ml sterile Murashige and Skoog (MS) media without or with carbon nanoparticle (CNP) in tissue culture containers. Further, seedlings were grown in short-day cycles (i.e., 8 h day and 16 h night) under white light for 7 d. The white light was obtained from Philips17 W F17T8/TL741 USA Alto II technology tubes fitted in plant growth chambers (Model-AR36, Percival, USA). Medium plates without CNP served as the controls. The intensity of incident white light was 80 µmol m^-2^ s^-1^. Seedling phenotype such as shoot length (SL), root length (RL), rootlet number (RLN) and nodal roots (NR) were recorded on 8^th^ day. Data presented the average of at least 2 biological replicates, and each biological replicate was a mean of at least 30 measurements.

#### Plant growth condition

For carbon nanoparticle (CNP) treatment in rice plants, growth conditions were adherent to as done in Panigrahy et al., (2021). Briefly, seedlings were grown on filter paper for 10 days, followed by growth on pooled pot up to one more month. The 40-day old seedlings were transferred to single plant per pot. CNP treatments (500 µg/ml) were added on the pot soil in every 15 days, i.e., on 45^th^, 60^th^, 75^th^, 90^th^ day.

### Abiotic stress treatments in rice seedlings

For imposing abiotic stress in seedling stage, rice seeds were de-husked and sterilized, followingly sown on 50 ml MS or CNP medium till 4 d before stress treatment. Cold and high temperature stresses were imposed at 18 °C and 35°C respectively. For PEG treatment, drought was induced with 10 % of PEG 6000 (Himedia, Cat #: GRM401, CAS #: 25322-68-3) at pH 5.7. For salt stress assay, 150 mM NaCl (Himedia, Cat #: MB023, CAS #: 7647-14-5) containing MS media were prepared with pH 5.7. For ABA treatment, 2-cis 4-trans-abscisic acid (Sigma, Cat # 862169) at 10µM was prepared in ethanol and sterile filtrated with 0.2 µm filter. It was mixed in the autoclaved medium just before pouring (lukewarm) in tissue culture containers. Five-d-old seedlings were carefully transferred on to medium containing PEG or salt or ABA plantons. For PEG, salt and high temperature, seedlings were grown for 7 more days before they were phenotyped for the growth parameters such as SL, RL, RLN on 13^th^ day. For cold treatment, seedlings were grown for one more day on MS medium (5d) before treatment for attaining higher growth to compensate the growth retardation due to cold. Hence, for cold and ABA treatment, phenotyping was done on 14^th^ day.

#### Cold or drought stress treatment in rice plants

Cold stress tolerance at 10 °C and drought stress tolerance was experimented in rice plants after four doses of CNP treatment in the reproductive stage. For imposing cold stress, top second leaves from 6 plants were incubated in water at 10 °C for nearly 10 d and observed for any leaf yellowing or deteriorating signs. Recovery from cold stress was done by keeping the cold-stressed leaves in room temperature for 24 h. Chlorophyll and carotenoid content was estimated from both the cold-stressed and recovery treated leaves. For drought treatment, water was withheld for 48 h and recovery assay was done 24 h after re-watering. Physiological analyses such as chlorophyll and carotenoid content, relative water content, malondialdehyde content, proline content, leaf temperature, relative humidity, leaf internal CO2 concentration, stomatal conductance, transpiration rate and vapor pressure deficit. were done from water-stresses leaf (top 2^nd^) samples in comparison with their respective control leaf samples. Data was obtained from 3 replicates each from 6 individual plants of untreated or CNP-treated plants.

### Water relation Parameters assessment

Leaf gaseous exchange and water relation parameters were measured using a CIRAS-3 Portable Photosynthesis System, Version 2.02, PP Systems, Amesbury, USA with parameter settings for PCL3 Universal Leaf Cuvette, 7 × 25 mm window. Settings for measurement were fixed with cuvette flow 300 CC min^-1^, analyser flow 100CC mm^-1^, H_2_0 reference 80%, temperature sensor measurement IR sensor, Leaf area 1.75 cm, boundary layer resistance 0.4 m^2^ s mol^-1^ and stomatal ratio 50%. Physiological parameters such as leaf temperature (Tleaf), relative humidity (RH), photosynthetically active radiation inside leaf (PARi), Assimilation (A_n_), stomatal conductance (gs), transpiration rate (E), vapour pressure density (VPD), internal carbon concentration (Ci) and water use efficiency (WUE) were recorded in control and CNP-treated plants in the middle part of top 2^nd^ leaf. The recordings were done before each CNP treatment and finally 15 days after (on the 125^th^ day) the last treatment. Hence, 45^th^, 60^th^, 75^th^, 90^th^, 125^th^ day measurements were noted as 0-day, 15^th^-day, 30^th^ -day, 45^th^ -day and 60^th^ - day of CNP-treatment respectively. Each data point is a mean of 108 recordings with 6 plants from each either untreated or CNP-treated plants. For each plant 18 recordings were done from 3 leaves in time series of 5 seconds interval, with 6 readings from each leaf.

### Tissue fixation and histological analysis using Confocal Microscopy

For stomata visualization in leaf tissue, samples from 9 d-old seedlings or 90 d-old plants (after 4^th^ CNP treatment) were fixed with FAA substituted with methanol (45% methanol, 10% formaldehyde, 5% Glacial acetic acid and 40% distilled water) under vacuum using a portable vacuum pump for 15-20 minutes. Leaf samples were stained with propidium iodide in (1:1000 v/v) phosphate buffer to visualise morphology of stomata. Tissues were mounted in glycerol. For visualization of CNP aggregates protocol from Allen et al (1940) with slight modification were followed. Briefly, seedling root tissues with or without CNP and / or nocodazole were fixed with FAA with vacuum overnight. On second day, samples were washed with ethanol, dipped in 0.1% eosin at 4°C. Next day, samples were washed with xylene wash series for 2 hrs, followed by moulding in paraplast at 65°C. On 8^th^ day, wax embedding was done at 4°C overnight. Sectioning was done and sample ribbons were kept in water-bath at 50 °C. Deparaffinization was done with xylene and ethanol. Mounting with glycerol was done after staining with safranin. A Leica SP8 confocal microscope system operating on a DMI600 inverted microscope, equipped with 63× NA 1.3 glycerol immersion lenses (Leica, Wetzlar, Germany), was used for imaging. High resolution Z stack images (1,024 pixels square) were recorded with 4-fold line averaging and a 1.00 μm step size. Along with concurrent transmitted light images, fluorescence images were collected from 420-500 nm using excitation at 405 nm. All image series were captured with the same imaging conditions. Images were modified using standard brightness and contrast settings in Photoshop (version CS4, Adobe Systems, San Jose, CA, USA) and ImageJ software.

### Physiological and Biochemical analyses of CNP treatment in rice seedlings

For all the physiological and biochemical assays, the 14^th^ d-old seedlings samples with or without cold treatment at 18 °C or 13^th^ d-old seedlings leaf samples with or without PEG treatment were used. All physiological and biochemical parameters were done thrice with three replicates in each set. Data were generated from the mean and the SEM. Significant differences were determined using one-way ANOVA using the Turkey’s multiple comparison in the Prism version 7.0 software. For Chlorophyll and carotenoid content, pigments were extracted in 80% acetone extract and estimated according to Arnon (1949) from the absorbances values at 663 nm, 647 nm and 480 nm (Panigrahy et al., 2018). Relative water content (RWC) was analysed using the formula, RWC = ((FW-DW) / (TW-DW)) × 100 Where: FW = Fresh weight; TW = turgid weight. Proline content was spectrophotometrically estimated at 520 nm according to Carillo et al. (2011), with little modification. Briefly, 20 mg leaf samples were extracted in 70% ethanol in 1:50 w/v proportion. Reaction mixture was prepared mixing 1% ninhydrin, 60% acetic acid, 20% ethanol. Leaf extracts were mixed with the reaction mixture (1:2 v/v), heated at 95°C for 10 min and supernatant was taken. For standard curve, L-proline (Hi-media, Cat # 147-85-3) was used. Malondialdehyde (MDA) content was determined according to Khan and Panda (2008) with little modifications. Briefly, 50 mg of leaf samples were ground with extraction buffer containing 0.25% thiobarbituric acid (TBA) and 10% trochloroacetic acid (TCA) followed by incubation at 95°C for 30 min. The extract was centrifuged at 10k for 10 min and absorbance of the supernatant was measured at 532 nm. Senthilkumar et al. (2021) For electrolyte leakage (EL) percentage, leaf samples were cut in to 0.3 cm × 1 cm pieces and prepared so as to have equal fresh weight with equal number of cuts. EL and were assayed according to Bajji et al. (2002) and (Flint et al. 1967)

#### Abscisic acid treatment

ABA at 10µM was prepared in ethanol and sterile filtrated. It was mixed in the autoclaved medium just before pouring (luke warm) in tissue culture containers. Dehusked and sterilized seeds were grown on MS medium till 4 d, after which they were trans with or without CNP and ABA separately.

### Endocytosis inhibitor treatment

Seedlings were grown in water for 8 days and nocodazole treatment was done according to Chem et al. (2018), briefly, 10µM in ethanol in stock was applied on the 3^rd^ day. Water uptake was estimated from the water left after 8 days. Water uptake/ seedling measured by subtracting seedling weight from the amount of water taken up. Root samples with or without inhibitor were processed for confocal microscopy.

#### Defence Hormone estimation in rice leaf samples

Hormone estimation was done in triplicates from ∼100 mg leaf samples of rice plants after 4^th^ CNP treatment on 90^th^ d. Estimation of defence hormones including abscisic acid (ABA), salicylic acid (SA) and jasmonic acid (JA) were done according to Vadassery et al., (2012). Leaf tissues were lypholized overnight after grinding with the help of a tissue lyser (Tissuelyser II, Retsch, Qiagen). Standards used were 1 ml of methanol containing 40 ng ml^-1^ of D6-jasmonic acid, (HPC Standards GmbH, Cunners dorf, Germany), 40 ng ml^-1^ D4-salicylic acid, 40 ng ml^-1^ D6-abscisic acid, and 8 ng ml^-1^ of jasmonic acid- [13C6] isoleucine conjugate for JA, SA, ABA and oxylipins respectively. The lipholyzed samples were extracted in methanol. Analysis of the samples was done with Exion LC (Sciex ®) UHPLC system using 0.05% formic acid as mobile phase A and acetonitrile as mobile phase B. for separation, Zorbax Eclipse XDB-C 18 column (50 x 4.6 mm, 1.8 µm, Agilent) was used coupled with a triple Quadruple-trap MS/MS system (Sciex 6500+) in negative ionization mode. The flow rate was 1 ml min^-1^ and isocratic elution profile were: up to 0.5 min: 5% B; 0.5 - 9.5 min: 5 - 42% B; 9.5 - 9.51 min: 42 - 100% B; 9.51 - 12 min: 100% B and 12.1 – 15 min: 5% B. Scheduled Multiple-reaction monitoring (MRM) is used to observe precisely analyte parent ion → product ion with detection window of 60 sec. Defense hormones were quantified relatively to the signal of their corresponding internal standard’s concentrations.

#### Abscisic acid quantification in rice seedling

Quantification of ABA from seedling samples was done according to Ali et al., (2011). Nearly 100 mg seedlings were homogenized with 80 % methanol at 4 °C. After filtering, samples were lyophilized with Heto DryWinner CT/DW 60E and redissolved in 0.5M phosphate buffer (pH 8). The samples were further purified through Sephadex G10 column and the eluted solution was lyophilized again. The residue was dissolved in acetonitrile and further filtered by 0.45 µm membrane filter (Waters) for HPLC analysis. The partially purified samples were analyzed using an HPLC system (Waters 1525 Binary HPLC) equipped with a photodiode-array (PDA) detector (Waters 2998) and RP-C18 column (5 mm packing; 250 x 4.6 mm internal diameter (ID). Acetonitrile with 0.5% acetic acid was used as a mobile phase and run isocratically with a flow rate of 0.3 ml min^-1^. The sample was injected with a SGE Analytical syringe (Melbourne, Australia) into the HPLC column through a Rheodyne (Oak Harbor, USA) injection valve equipped with 100 ul sample loop. The detection wavelength was 254 nm. Standard stock solution of pure ABA was prepared by dissolving 1 mg of ABA (Sigma, extra pure) in 1 ml of HPLC grade acetonitrile. Quantification of ABA in the rice seedlings was carried out by comparing the standard curve generated through Breeze 2 software of Waters HPLC.

### RNA isolation and genome-wide transcript expression analysis

#### RNA isolation

Total RNA was isolated from rice flag leaves and *Arabidopsis* seedlings for genome-wide microarray analysis and transcriptome analysis respectively. For microarray analysis, rice flag leaves from untreated or CNP-treated plants were sampled after 4^th^ CNP treatment on the day of complete emergence of the panicle from the main stem. For transcriptome sequencing, 14 d-old *Arabidopsis thaliana* (*At*) Col-0 seedlings grown of MS or CNP medium were harvested. All samples were sampled with liquid N_2_ and Experiments were performed in duplicates. RNA was extracted from ∼200 mg frozen rice leaves or 1 g of *At* seedlings using Qiagen RNAeasy Plant Mini kit (Cat#74904). Briefly, leaves were homogenized using TOMY smasher homogenizer with liquid N_2_ and RLT buffer. Lysate was centrifuged and supernatant was mixed with RW1 buffer and loaded onto Qiagen RNeasy column and further steps including on-column DNase treatment (Qiagen RNase free DNase Cat#79254) were followed as per manufacturer’s guidelines. RNA was eluted in Nuclease free water (Cat#AM9938, Ambion, USA). Quantity and purity of the RNA were evaluated using the Nanodrop 2000 Spectrophotometer (ThermoFischer, USA) from 260/280 values. Integrity of RNA was verified on the 2100 Bioanalyzer (Agilent, USA) using rRNA 28S/18S ratios and RNA integrity number (RIN).

#### Microarray experimentation and data analysis

The rice RNA samples for gene expression were labelled using Agilent Quick-Amp labelling Kit (p/n5190-0442). The total RNA was reverse transcribed at 40°C using oligo dT primer tagged to a T7 polymerase promoter and converted to double stranded cDNA. This cDNA was used as template for cRNA generation using in vitro transcription and incorporation of the dye Cy3 CTP(Agilent). Cleaning of cRNA was done using Qiagen RNeasy columns (Qiagen, Cat #: 74106) and quality assessed for yields and specific activity using the Nanodrop ND-1000. Fragmentation (at 60 °C) and hybridization (at 65° C for 16 hours) were done using the Gene Expression Hybridization kit of (Agilent Technologies, In situ Hybridization kit, Part # 5190-0404). An Agilent Rice Gene Expression Microarray 8x60k (AMADID 64722) was used for hybridization. Hybridization was carried out in Agilent’s Surehyb Chambers. Washing using Agilent Gene Expression wash buffers (Agilent Technologies, Part # 5188-5327) and scanning was done using the Agilent Microarray Scanner (Agilent Technologies, Part Number G2600D). Microarray data analysis was performed from the extracted raw data and was analyzed using Agilent GeneSpring GX (Version14.5) software. Normalization was done in GeneSpring GX using the 75^th^ percentile shift method was adapted to obtain the fold change (FC). Significantly down or up regulation (differentially expressed genes, DEG) were identified with a cut off range in the logbase_2_ of FC at ≤ -0.6 and ≥ 0.6 respectively in the test samples with respect to control sample. Statistical student T-test p-value among the replicates was calculated based on volcano Plot Algorithm. Differentially regulated genes were clustered using hierarchical clustering based on Pearson coefficient correlation algorithm to identify significant gene expression patterns. For function and pathway analysis differentially expressed genes were validated using Biological Analysis tool DAVID (http://david.abcc.ncifcrf.gov/).

#### Transcriptome sequencing and data analysis

*Arabidopsis thaliana* RNA sequencing libraries were prepared with Illumina-compatible NEBNext® Ultra™ II Directional RNA Library Preparation Kit (New England BioLabs, MA, USA). Five hundred ng of total RNA was taken for mRNA isolation. Fragmented and primed mRNA was further subjected to first strand synthesis followed by second strand synthesis. The double stranded cDNA was purified using JetSeq magnetic beads (Bio, # 68031). Purified cDNA was end-repaired on Illumina HiSeq X Ten sequencer (Illumina, San Diego, USA), adenylated and ligated to Illumina adapters as per NEBNext® Ultra™ II. Sequencing libraries were purified with JetSeq Beads. Illumina-compatible sequencing libraries were quantified by Qubit fluorometer (Thermo Fisher Scientific, MA, USA) and fragment size distribution was analyzed on Agilent 2200 TapeStation. The Illumina libraries showed average fragment size range of 433 bp which was sufficient concentration to get the sequencing data. Transcriptome analysis was performed by processing the raw data for removal of low-quality data and adapter sequences. The data obtained from the sequencing run was demultiplexed using Bcl2fastq software v2.20 and FastQ files were generated based on the unique dual barcode sequences. The raw reads were processed using FastQC2 (v0.11.8) software for quality assessment and pre-processing which includes removing the adapter sequences and low-quality bases (<q30) using TrimGalore3.

Illumina sequencing generated a total of 65713,399 paired end raw data. Nearly 98.08% of total reads were retained as high quality (>Q30) data. The pre-processed 16.1 million high-quality reads were considered for alignment with reference *Arabidopsis thaliana* genome using a splice aligner using Hisat24 with the default parameters to identify the alignment percentage and classified into aligned and unaligned reads. About 97.58% of the reads were aligned to the reference genome. HTSeq5 was used to estimate and calculate gene abundance. DESeq6 used the absolute counts for genes to identify and used for differential expression calculations. DEG were categorized as Up, Down based on cut-off of log_2_fold change +/-1.0 value respectively. Genes were annotated using homology approach to assign functional annotation using BLAST7 tool against “Viridiplantae” data from the Uniprot database. The DEG were analysed for GO annotation and pathway analysis to understand the functional role of each expressed gene. A match was found when e-value less than e^-5^ and minimum similarity greater than 30%. Pathway analysis was done by using KAAS8 Server. Enrichment analysis for all up and down regulated genes was performed using ShinyGO9 tool.

#### Real-time PCR (Q-PCR) analysis

For real-time PCR analysis, plants tissues (rice flag leaves and *Arabidopsis* seedlings) were sampled with liquid N_2_. The growth condition and sampling procedure of rice flag leaves for this purpose was similar to as that for the microarray expression analysis (section). *Arabidopsis* seedlings (10 d-old) were sampled after 12 h of onset of light in the long-day cycle (Zt-12). RNA was extracted from using RNeasy Plant Mini Kit Qiagen (Cat #74104) and one µg of total RNA was used to obtain cDNA using Reverse Transcription Super-mix (Cat #1708840). Quantitative RT-PCR (qRT-PCR) was carried out using the Bio-Rad laboratories’ CFX384 Touch^TM^ Real time detection system and iTaq^TM^ universal SYBR green super mix following the manufacturer’s instructions as done previously (Kumar et al., 2018). Primer Quest tool (Integrated DNA Technologies, Inc., USA) was used to design gene-specific primers (Table S3). All reactions were carried out in hard shell 384-well PCR plates (Bio-Rad, Cat #: HSP3805). ACTIN was used to normalise transcript levels. The qRT-PCR reactions were done in triplicates from two biological replicates. Data represents as the mean ±SEM. The relative quantification of genes was analyzed using standard 2^-ΔΔct^ as described by Pfaffl (2001). For relative fold-change, the respective expression value for the same gene from untreated plant leaf samples was taken as 1.

## Acknowledgement

This work was funded by Department of Science and Technology (DST) Women Scientist-A (WOS-A) grant, Government of India, File No: SR/LS/WOS-A/369/2018. Central Infrastructural Facility and technical support of NISER, Odisha is highly acknowledged. Institutional support from SOA University is acknowledged. The transcriptome and microarray genomics experiments were performed at Genotypic Technology, Bangalore. Abscisic acid quantification for seedlings was done at Visva-Bharati Central University), West Bengal. Abscisic acid quantification for flag leaves was done at Plant metabolite service facility, National Institute of Plant Genome Research (NIPGR), New Delhi.

## Authorship contribution statement

MP has designed the project, acquired funds. LM and JPK have done the experimental work. AK has done real-time PCR experiments. SM and JR have done the ABA content determination. GM has done the microscopy. KCP and GRR have acquired funds, provided infrastructural and institutional support, helped in designing, trouble shooting. All authors have read and agreed to the MS.

## Data Availability

Microarray data sets for CNP treatment in rice plants can be accessed at: NCBI GEO Accession No: GSE200357, Title: “Effect of Carbon Nanoparticle Treatment in Rice Adult plants”, https://www.ncbi.nlm.nih.gov/geo/query/acc.cgi?acc=GSE200357

Transcriptome sequencing data sets for CNP treatment in Arabidopsis seedlings can be accessed at NCBI SRA Submission Portal with the reviewer’s link https://dataview.ncbi.nlm.nih.gov/object/PRJNA825704?reviewer=kefmtv4hfnnu7siae3kch7uekf

## Supporting Information

**Figure S1.**
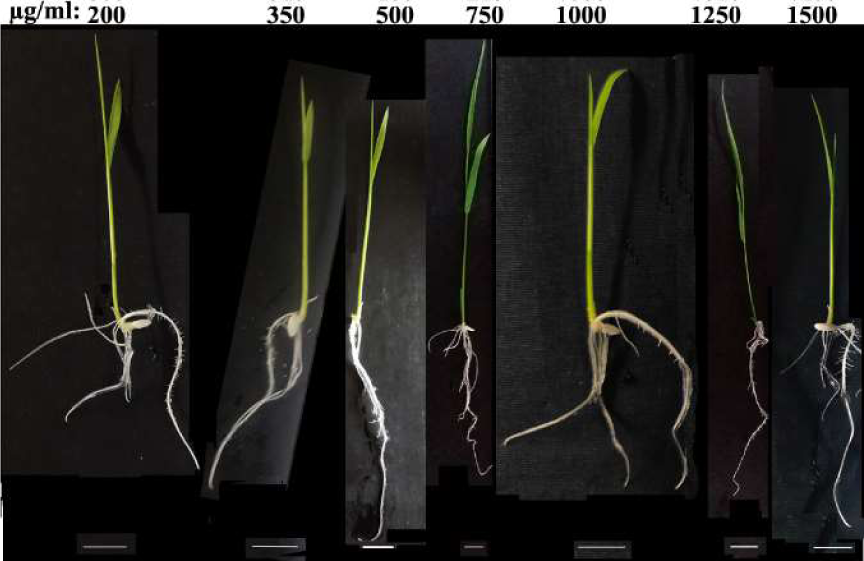
Representative seedling pictures of dosage dependence effect of Carbon nanoparticle. The concentration of CNP in µg ml^-1^ is indicated on the top of each seedlings. Scale bar: 2 cm

**Figure S2.**
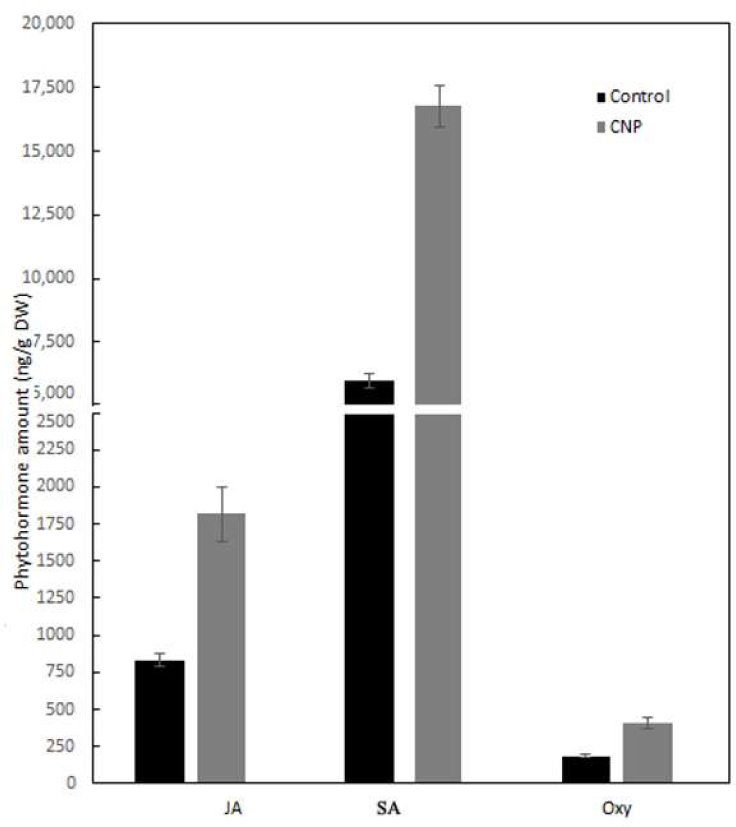
Endogenous defence hormone quantification in CNP treated seedlings. Defence hormones including Jasmonic acid (JA), Salicylic acid (SA) and Oxylipins (Oxy) were quantified in adult leaf samples after 4^th^ CNP treatment. Top second leaf was used for lyophilization and processed as explained in the section 2.6.1. Data represented is a mean of three replicative measurements. Statistical significance was obtained from one-way ANOVA using the Turkey’s multiple comparison in the Prism version 7.0 software, and were represented as *, *P* ≤ 0.05, **, *P* ≤ 0.01 and ***, *P* ≤ 0.001.

**Figure S3.**
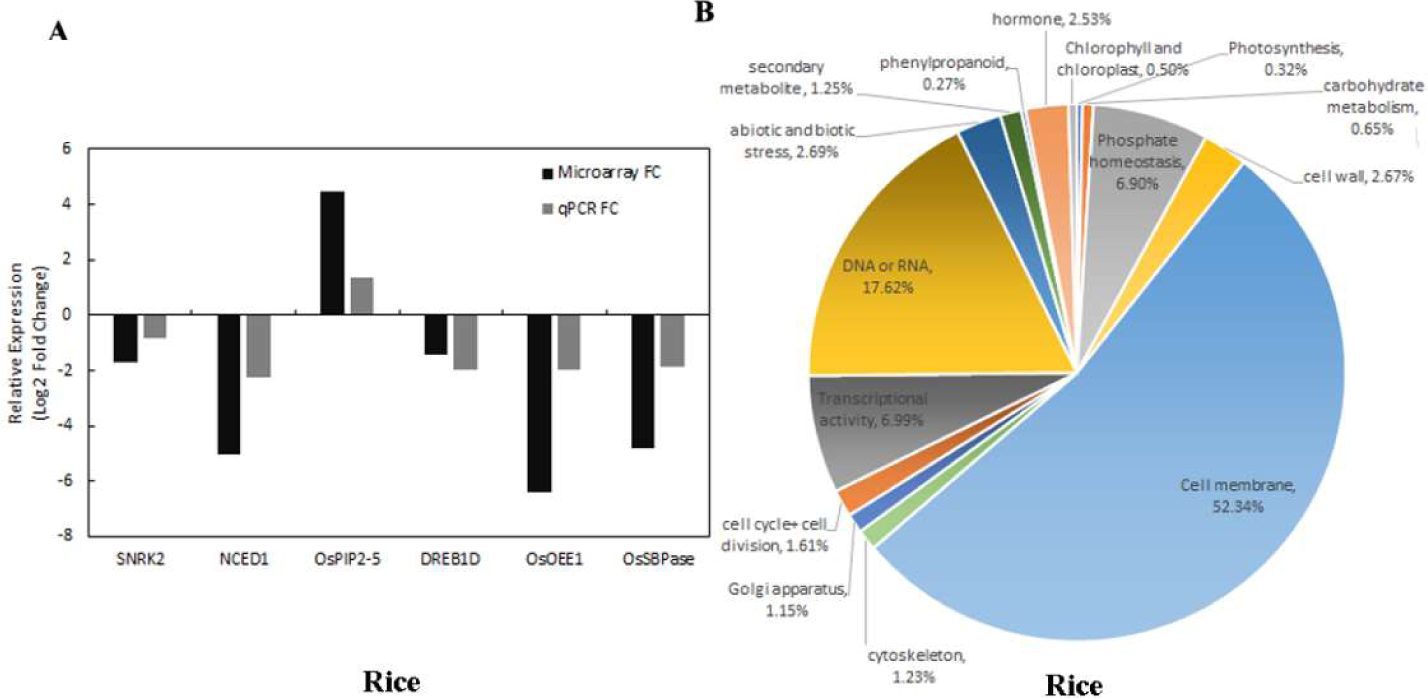
Function & pathway analysis and validation of microarray expression data from CNP-treated and control leaf samples. (A): Correlation plot among replicates of control and CNP-treatment samples (B) Differentially expressed genes (DEGs) obtained from microarray expression analysis were categorized according to their transcript details obtained from Rice Annotation Project (RAP-DB), MSU and CGSNL Databases. Functional details were obtained from GO annotations, KEGG database, UNIPROT and Oryzabase database. Pathway categorization was done according to the biological process, cellular category and molecular function. Percentage was obtained by calculating from total number of DEGs. (C): collinearity of the Log_2_ fold change values obtained from microarray expression analysis and qPCR.

**Figure S4.**
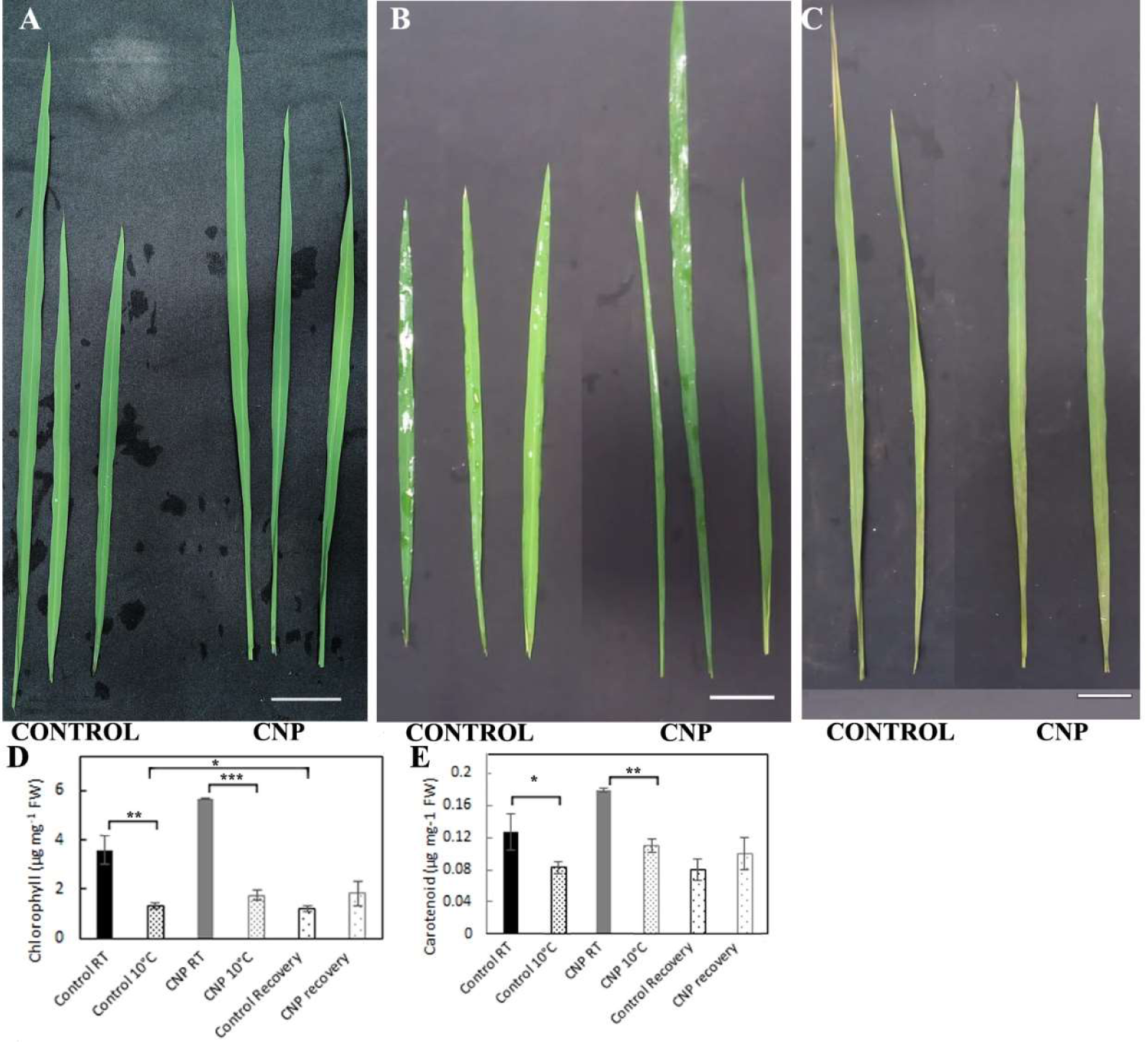
Cold treatment assay in flag leaves of CNP-treated and control plants. Cold stress was imposed in top second leaves taken from rice plants in the reproductive stage (after 4^th^ CNP treatment). They were floated in equal amount of water and incubated in white light at 10°C for 10 days. (A) Leaves before cold stress (B) Leaves after cold stress (C) Leaves from cold stress were incubated for 24 h recovery at room temperature. (D) chlorophyll (E) carotenoid content were estimated from control and CNP-treated leaves mentioned above. The experiment was done thrice from leaves of 3 biological replicates. Statistical significance was obtained from one-way ANOVA using the Turkey’s multiple comparison in the Prism version 7.0 software, and were represented as *, *P* ≤ 0.05, **, *P* ≤ 0.01 and ***, *P* ≤ 0.001.

**Table S1.**
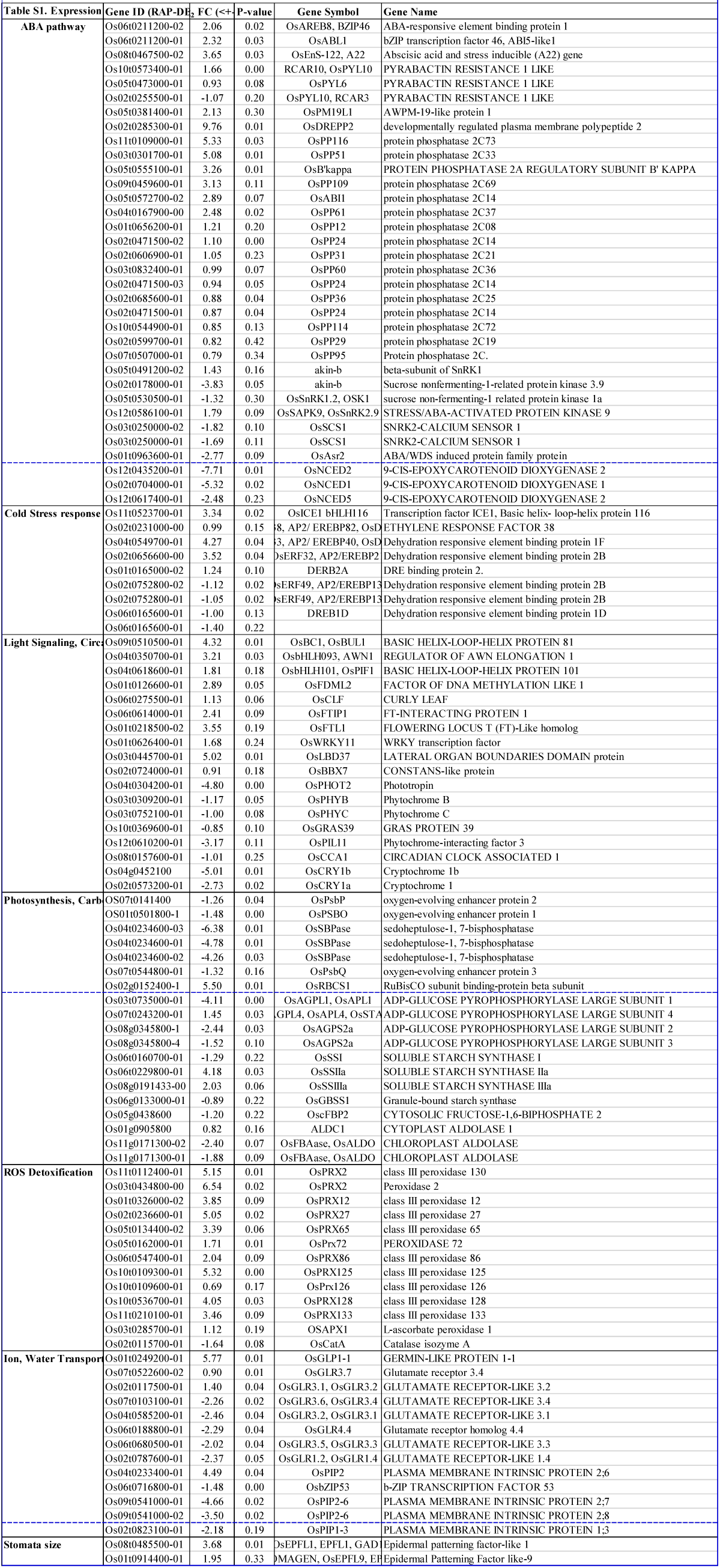
Expression level of Transcripts of some selected pathway from microarray analysis in rice. Flag leaves on the day of final emergence of the panicle from the CNP treated and untreated plant were sampled with liquid N2 after 4^th^ CNP treatment. Microarray analysis was done from two biological replicates each category. Normalization and Fold change was calculated as described in the materials and methods section. Transcript details are taken from Rice Annotation Project Database (RAP-DB). Pathway analysis was done using GO Ontology, KEGG and Biological Analysis tool DAVID (http://david.abcc.ncifcrf.gov/)

**Table S2.**
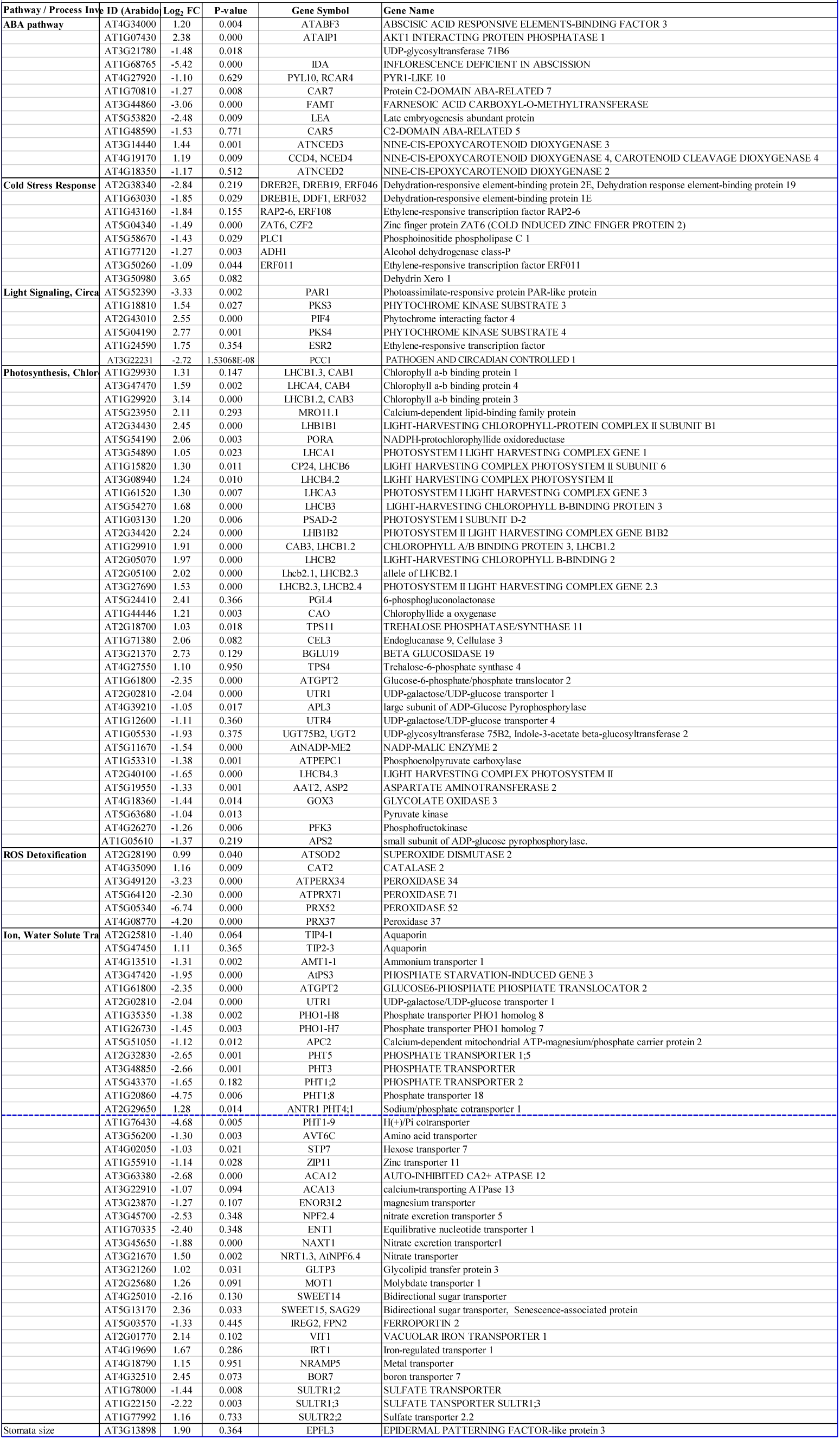
Expression level of Transcripts of some selected pathway from transcriptome sequencing analysis in Arabidopsis thaliana. 16-day old CNP seedlings from two biological replicates each category of treated and untreated were sampled with liquid N2 for transcriptome sequencing. Normalization and Fold change was calculated as described in the materials and methods section. Transcript details are taken from TAIR database.

**Table S3. Primers used in this study**

